# Lipid Droplets Fuel Small Extracellular Vesicle Biogenesis

**DOI:** 10.1101/2022.10.24.513202

**Authors:** Géraldine C. Genard, Luca Tirinato, Francesca Pagliari, Jessica Da Silva, Alessandro Giammona, Fatema Alquraish, Marie Bordas, Maria Grazia Marafioti, Simone Di Franco, Jeannette Janssen, Daniel Garcia-Calderón, Rachel Hanley, Clelia Nistico, Yoshinori Fukasawa, Torsten Müller, Jeroen Krijgsveld, Matilde Todaro, Francesco Saverio Costanzo, Giorgio Stassi, Michelle Nessling, Karsten Richter, Kendra K. Maass, Carlo Liberale, Joao Seco

## Abstract

Despite an increasing gain of knowledge regarding small extracellular vesicle (sEV) composition and functions in cell-cell communication, the mechanism behind their biogenesis remains unclear. Here, we revealed for the first time that the sEV biogenesis and release into the microenvironment are tightly connected with another important organelle: Lipid Droplets (LD). We have observed this correlation using different human cancer cell lines as well as patient-derived colorectal cancer stem cells (CR-CSCs). Our results showed that the use of external stimuli such as radiation, pH, hypoxia, or lipid interfering drugs, known to affect the LD content, had a similar effect in terms of sEV secretion. Additional validations were brought using multiple omics data, at the mRNA and protein levels. Altogether, the possibility to fine-tune sEV biogenesis by targeting LDs, could have a massive impact on the amount, the cargos and the properties of those sEVs, paving the way for new clinical perspectives.

**Significance Statement:** 

## Introduction

In 2013 Professors James E. Rothman, Randy W. Schekman and Thomas C. Südhof were awarded with the Nobel Prize for their discoveries of machinery regulating vesicle traffic, a major transport system in human cells (1). Their and other groups’ works highlighted the importance of intra- and extracellular vesicles (EVs) in the cell-cell communication and their ability to modulate the cellular microenvironment.

Almost all mammalian cells produce EVs, defined as “lipid bilayer-enclosed extracellular structures” of different size and intracellular origin. EVs are characterized by their size, cell origin, molecular composition, and functions (2). Small extracellular vesicles (sEVs) are distinguished from other EV subtypes by their small size (30 - 200 nm) and their ability to travel along the blood and lymph streams to reach distant organs from their sites of origins. Since they carry intracellular content of donor cells (including DNA, RNA, proteins, and lipids), those sEVs influence the fate of acceptor cells (3, 4). Their roles have been described in many physiological and pathological conditions, such as cancer, cardiovascular disease, immune response, and regeneration (5). In a tumor context, cancer cell–derived sEVs are believed to be secreted in large amount, with the ability to remodulate the tumor microenvironment and the tumor progression through various mechanisms, including immune evasion(6), proliferation, invasion, or metastasis (5).

sEVs have two different subcellular origins, either endosomal or non-endosomal, making them heterogenous. In particular, sEVs of endosomal origin, so-called *exosomes*, are nanoparticles released through the fusion of multivesicular bodies (MVBs) (containing intraluminal vesicles (ILVs)) with the plasma membrane (2). The non-endosomal pathway generates sEVs devoid of CD63, CD81 and CD9 or sEVs enriched in ECM and serum-derived factors (7).

As all sEVs are shaped by lipids, we hypothesized that a potential common source builds up the surrounding membrane: either coming from the recycling of plasma membrane within the endosomal pathway or through a new source of phospholipids.

Lipid Droplets (LDs) have been considered as mere fat storage organelles for a long time, although important evidence could be traced back to the early 1960’s (8). As of today, LDs are well recognized as fundamental cellular hubs involved in many physiological as well as pathological processes, including cancer (9, 10). Nevertheless, many open questions about their formation, composition and role remain to be fully elucidated.

LDs are spherical organelles, which are found in the cytoplasm, and in some cases, in the nucleus of all eukaryotic cells(11). They are characterized by a lipid-rich core (cholesterol esters (CEs) and triacylglycerols (TAGs)) surrounded by a phospholipid monolayer (12). Although the LD-protein repertoire is cell-specific and influenced by the methodology used for their isolation, to date, more than 150 specific LD-proteins have been detected in mammalian cells (13).

In addition to their role in membrane biosynthesis, LDs are very active organelles due to their continuous cycle of growth and consumption reflecting the cell status needs (13). In this regard, during cell expansion and division (which require membrane enlargement and increased biosynthesis of phospholipids), the fatty acids stored as TAGs in the LD core are mobilized either by lipolysis or by lipophagy (13). This allows the cell to sustain several metabolic processes and membrane biosynthesis.

LDs were associated with numerous other functions. For example, LD accumulation protects cells from oxidative stress damage by sequestering free fatty acid (14). In the same context, LD increase is considered as a cancer stem cell marker in many tumors (15, 16) and as a cell signature for radioresistance (17). Moreover, a role for LDs in the immune system modulation has been also reported in colorectal cancer (18).

To carry out their multiple roles, LDs need to “interact” with other cellular players. To do so, they establish physical contact with several organelles, like the endoplasmic reticulum (ER), peroxisomes, lysosomes, mitochondria, and endosomes (13).

Several reports suggested a connection between the lipid incorporation into LDs and the intracellular vesicle formation (13, 19). Interestingly, it was seen that the adipose tissue, whose cells contain the largest amount of LDs, is responsible for the highest number of secreted sEVs (AdExos)(20). It was also shown that these lipid-filled AdExos are then used by macrophages as a source of lipids (21).

Based on this evidence, we decided to investigate the potential connection between LDs and sEVs. To this purpose, we used different commercial human cancer cell lines (colon, lung, pancreatic and breast cancer cells) as well as patient-derived CR-CSCs. By using several means, we analyzed the impact of modulating the LD content on sEVs and the connection LDs – sEVs. Indeed, we adopted different external stimuli (such as distinctive pH, oxygen concentration, and ionizing radiations) or used LD inhibitors and silencing of Ferritin Heavy Chain 1 (FTH1), since its role in the LD formation has been already shown (17).

## Results

### The number of LDs strongly correlates with the release of sEVs in colorectal cancer cell lines

To evaluate if there is a possible connection between cellular LD content and sEV release, we first compared both the number of LDs and the average amount of released sEVs per cell, in two different colorectal cancer cell lines, LoVo and HT29 (**Fig 1**). As shown by z-stack projections of confocal microscopy images and by the associated LD quantification, HT29 contained significantly more LDs per cell than LoVo 72h after seeding (**Fig 1A**). In parallel, the released sEVs were studied for both cell lines 72 hrs after seeding. The sEV isolation protocol was used as described in (22) and pictured in **Fig S1A**. The purity of the sEV samples was validated by observing the presence of exosomal markers (CD81, Tsg101 and CD63) as well as the absence of Golgi (GM130), endoplasmic reticulum (Calnexin), mitochondrial (Cytochrome C) and plasma membrane and cytoplasmic (Enolase 1) markers in the sEV preparations. In accordance with the literature (23), we found the presence of Hsc-70 both in the cellular and sEV fractions, with a predominance for the cellular fraction (**Fig S1B**). As expected, EM analysis of the sEV preparations showed a size ranging from 30 nm to 200 nm for the isolated sEVs (**Fig 1B**). Similarly, we could determine the number of particles and their size by using Nanoparticle Tracking Analysis (NTA). The average size of particles peaked at 148 nm for LoVo cells and 133 nm for HT29 cells (**Fig S1C**). The NTA measurement (**Fig 1C**) and the protein quantification (**Fig S1D and S1E**) also confirmed a higher amount of sEVs released per cell for HT29 as compared to LoVo cells. We next aimed to identify exosomal markers by western blotting to confirm the higher number of sEVs released by HT29 cell line. By loading the same volume of each sample, we observed that exosomal markers (CD9, CD63, CD81 and hsc-70) were significantly more expressed in the sEV fractions collected from HT29 than LoVo cell line (**Fig 1D**). Finally, as the number of LDs might be heterogeneous among the same cell line, we sorted HT29 cells based on their LD content.

**Figure 1.**
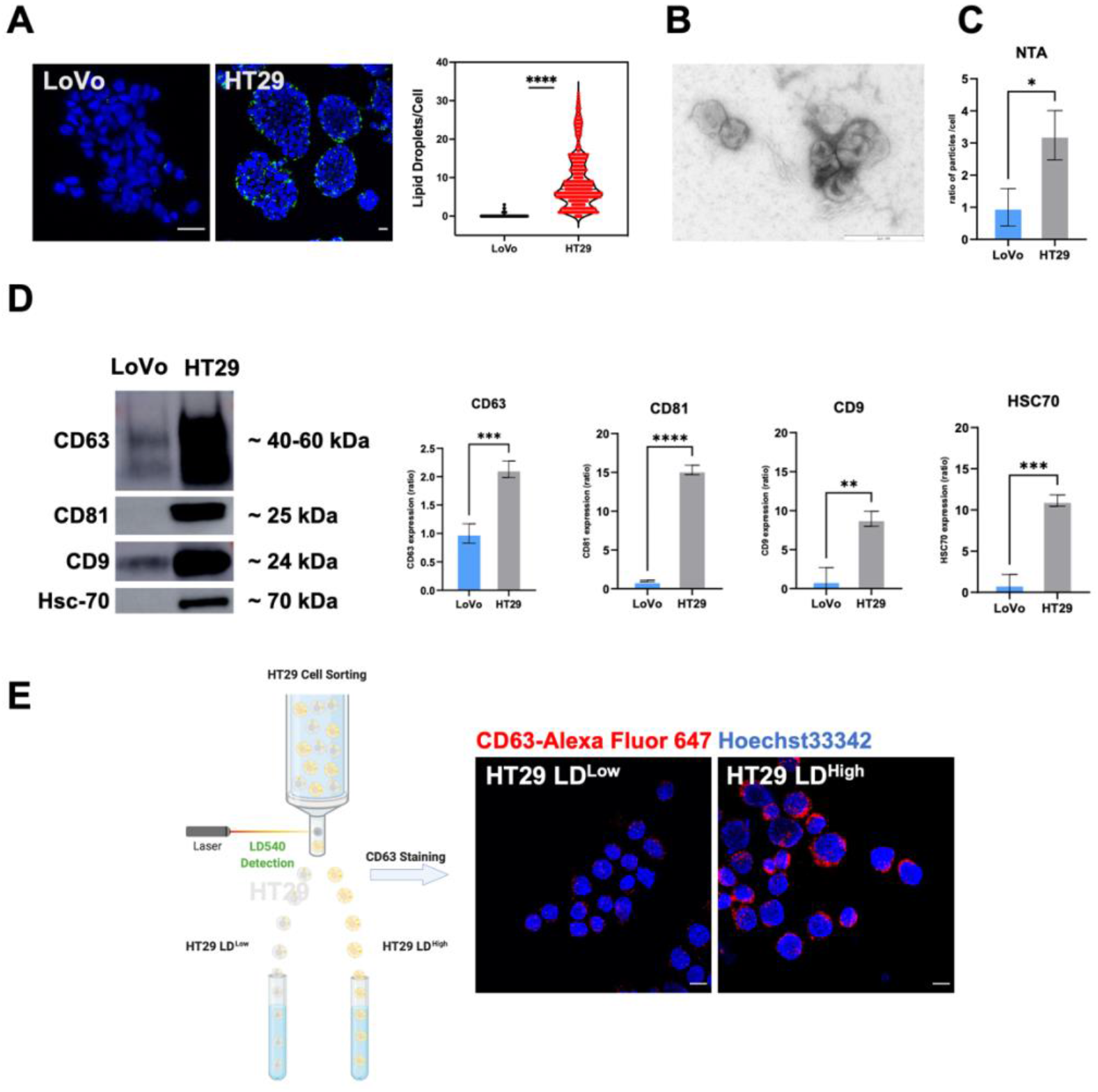
Analysis of LD content and sEV release in LoVo and HT29 colorectal cancer cell lines. **A**) HT29 and LoVo cell lines were stained with LD540 (green) for LDs and DAPI (blue) for nuclei and imaged at the confocal microscope with a 100X objective (Leica Microsystems; Concord, Ontario, Canada). The pinhole was set for a slice thickness of 17.4 μm, with an interval between slices of 0.9 μm. Z-projection of the z-stack acquisitions is shown (left). Displayed are the merged images of the LD540 and DAPI staining from one independent experiment (Scale bar, 20 μm). The graph represents the changes in LD content for LoVo and HT29 cell lines. Images were analyzed using ImageJ for mean of LDs per cell. Comparisons between groups are shown with corresponding p-values (unpaired Student’s t-test). Error bars represent the means ± SD. n=1 (LoVo: N = 532 cells; HT29: N = 4645 cells). **B**) High-resolution transmission electron micrograph of sEVs isolated from HT29 media taken with Zeiss EM 910 at 100 kV. Uranyl acetate negative staining reveals that purified sEVs have a cup-shaped morphology enclosed by a lipid bilayer. The diameter of sEVs is around 90–100 nm. The presented image has a magnification of 16000 x in TEM mode. The size bars on the image represent 250 nm. **C**) Ratio of particle number per cell for the sEV fractions (F2) released by LoVo and HT29 by nanoparticle tracking analysis (NTA). Comparisons between groups are shown with corresponding p-value. Unpaired students t-test was performed. Error bars represent the means ± SD from three independent experiments. **D**) Western blot for the sEV pellets (100K) obtained by differential ultracentrifugation combined with SEC for LoVo and HT29 cells. The same sample volume (19.5 μL) was loaded onto the 10% acrylamide gel. The results presented here are representative of three independent experiments. The intensity of the bands corresponding to HT29 proteins was normalized by the intensity of the LoVo proteins band. Unpaired students t-test was performed. Error bars represent the means ± SD from three independent experiments. **E)** HT29 cells were stained with LD540 for LDs and sorted based on their 10% brightest and and 10% dimmest LD540 fluorescence values. Thereafter, sorted HT29 cells were spun on slides using cytospin and were directly fixed, permeabilized and stained for CD63 (MVBs) and DAPI (nuclei). Cells were then imaged at the confocal microscope with a 100X objective (Leica Microsystems; Concord, Ontario, Canada). Displayed are the merged images of the CD63 and DAPI stainings (Scale bar, 20 μm). * ≤ 0.05; ** ≤ 0.01; *** ≤ 0.001 and ****≤ 0.0001.

Thereafter, the multivesicular bodies (MVBs) were assessed by confocal microscopy. The images indicated a high MVB numbers for the HT29 LD^High^ fraction in comparison to the HT29 LD^Low^ counterpart (**Fig 1E**). Altogether, these results suggest that the intracellular LD content followed the same trend as the released sEVs.

### Inhibition of LD metabolism reduces sEV release

Thereafter, we decided to target LD biosynthesis in HT29 cells by using two lipid inhibitors affecting two different steps of the LD biogenesis (**Fig 2A**).

**Figure 2.**
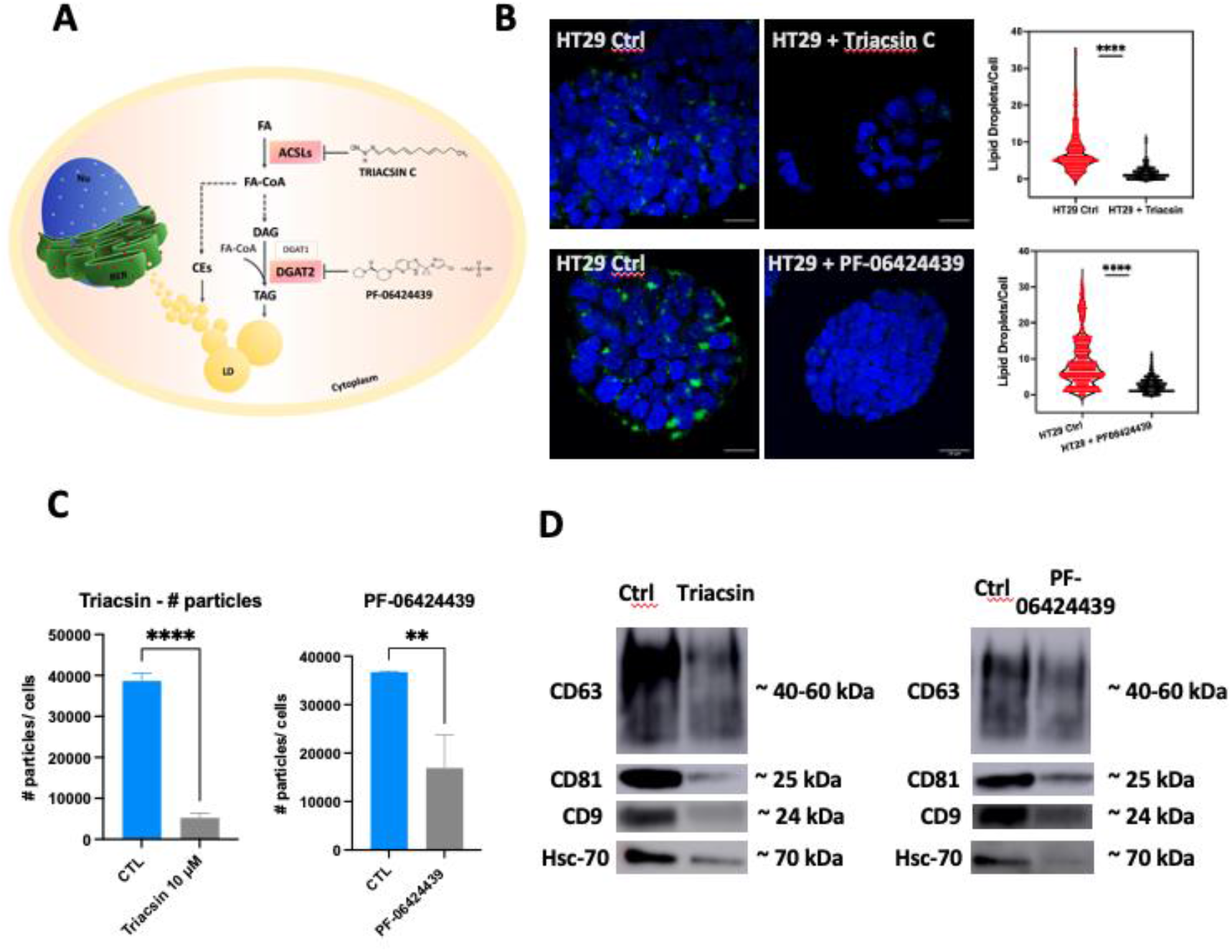
LD content inhibition reduces sEV release. **A**) Representation of the mechanism of action for Triacsin C and PF-06424439. **B**) HT29 cells control or treated, either with 10 μM Triacsin C or 30 μM PF-06424439 for 72 hrs, were stained with LD540 (green) for LDs and DAPI (blue) for nuclei and imaged at the confocal microscope with a 100X objective (Leica Microsystems; Concord, Ontario, Canada). The pinhole was set for a slice thickness of 17.4 μm, with an interval between slices of 0.9 μm. Z-projection of the z-stack acquisitions is shown (left). The merged images of the LD540 and DAPI staining from one independent experiment are displayed (Scale bar, 20 μm). The graph represents the changes in LD content for HT29 cell line treated or not with one of the two inhibitors used in this experiment. Images were analyzed using ImageJ for mean LDs per cell. Comparisons between groups are shown with corresponding p-values (unpaired Student’s t-test). Error bars represent the means ± SD. n=1 (HT29 CTL Triacsin C: N = 575 cells; HT29 treated with Triacsin C: N = 430 cells; HT29 CTL PF: N = 860 cells; HT29 treated with PF-06424439: N = 1083 cells). **C**) Ratio of particle number per cell for sEV factions (F2) released by HT29 control or treated with LD inhibitors using NTA. Unpaired students t-test was performed. Error bars represent the means ± SD from three independent experiments. **D**) Western blot for the sEVs pellets (100K) obtained by differential ultracentrifugation combined with SEC for HT29 cells control or treated, either with Triacsin C 10μM or PF-06424439 30 μM. The same sample volume (19.5 μL) was loaded onto the 10% acrylamide gel. The results presented here are representative of three independent experiments. * ≤ 0.05; ** ≤ 0.01; *** ≤ 0.001 and ****≤ 0.0001, ns = not significant.

The first drug acts as an inhibitor of long fatty acetyl-CoA synthetases (Triacsin C), while the second one blocks the glycerolipid synthesis (PF-06424439). Triacsin C and PF-06424439 were used at a concentration of 10 μM and 30 μM respectively. The choice of the inhibitor concentrations was made based on the literature for Triacsin C (18) and on the evaluation of LD and sEV numbers per cell for PF-06424439. Both inhibitors induced a cellular LD number reduction 72 hrs after incubation, as shown by confocal analysis and the associated quantification (**Fig 2B**). The LD decrease was correlated to a drop of sEV released in the supernatant by HT29 cells (**Fig 2C**) and to a reduction of the protein concentration within the sEV fraction (**Fig S2A**). In addition, a lower protein expression of exosomal markers (CD9, CD63, CD81 and hsc-70) as shown in **Fig 2D,** was observed 72 hrs after incubation with both inhibitors (**Fig 2D**). A quantification of exosomal marker expression was performed on 3 independent experiments emphasizing the difference between the control and the treated conditions (**Fig S2B**).

Altogether, those results strengthen the connection between LDs and sEVs.

### Iron metabolism supports the connection between LDs and sEVs

It is now quite well established that there is an interplay between the iron and the lipid metabolisms. In a previous work (17), we demonstrated that Ferritin Heavy chain (FTH1) – a key enzyme involved in cytoplasmic iron storage and redox homeostasis – regulated the cellular LD content. Therefore, we thought to use the same experimental system, based on short hairpin RNA targeting FTH1 (shFTH1) or scrambled RNA (shRNA) in the MCF7 cell line, to evaluate the sEV biogenesis. First, we collected proteins from MCF7 shRNA and MCF7 shFTH1 to conduct a full proteome analysis. From this analysis, 543 proteins were found to be upregulated (Log2 Fold change > 1.2) and 770 proteins downregulated (Log2Fold < 0.833) in MCF7 shFTH1 cells (**Fig 3A)**. We then confirmed that metabolic pathways, including small molecule metabolic processes and cellular catabolic processes, were downregulated (**Fig S3A**) in MCF7 shFTH1 cells. In addition, the expression of proteins involved in adipogenesis, fatty acid metabolism as well as lipoprotein and cholesterol synthesis was mostly downregulated in MCF7 shFTH1 cells (**Fig S3B**). In particular, 31 proteins involved in the lipid metabolism were upregulated while 46 proteins were downregulated in the MCF7 shFTH1 cell line. Using String and Cytoscape software, we found that the “extracellular vesicle” pathway was downregulated, among others, in MCF7 shFTH1 cells (**Fig 3B**). A closer look to the exosomal pathway highlighted that 62.7% of proteins related to the exosomal pathway were downregulated in MCF7 shFTH1 cells as compared to the MCF7 shRNA ones (**Fig 3C**). In accordance with these results, NTA analysis emphasized fewer sEVs/cell released from MCF7 FTH1 cells as compared to MCF7 shRNA cells (**Fig 3D**). By analyzing the protein expression of exosomal markers (Annexin V, Flotillin-1, CD81 and CD9) on the same sEV sample volume, we evidenced a lower expression of those markers in MCF7 shFTH1 than in MCF7 shRNA cells (**Fig 3E**). The proteomic results strengthen this outcome, as the expression of almost all exosomal markers was downregulated in MCF7 shFTH1 cells as compared to MCF7 shRNA ones (**Fig 3F**). Altogether, these results confirmed that sEV amount is directly correlated to the cellular LD content and that iron metabolism is upstream from the LD-sEV connection.

**Figure 3.**
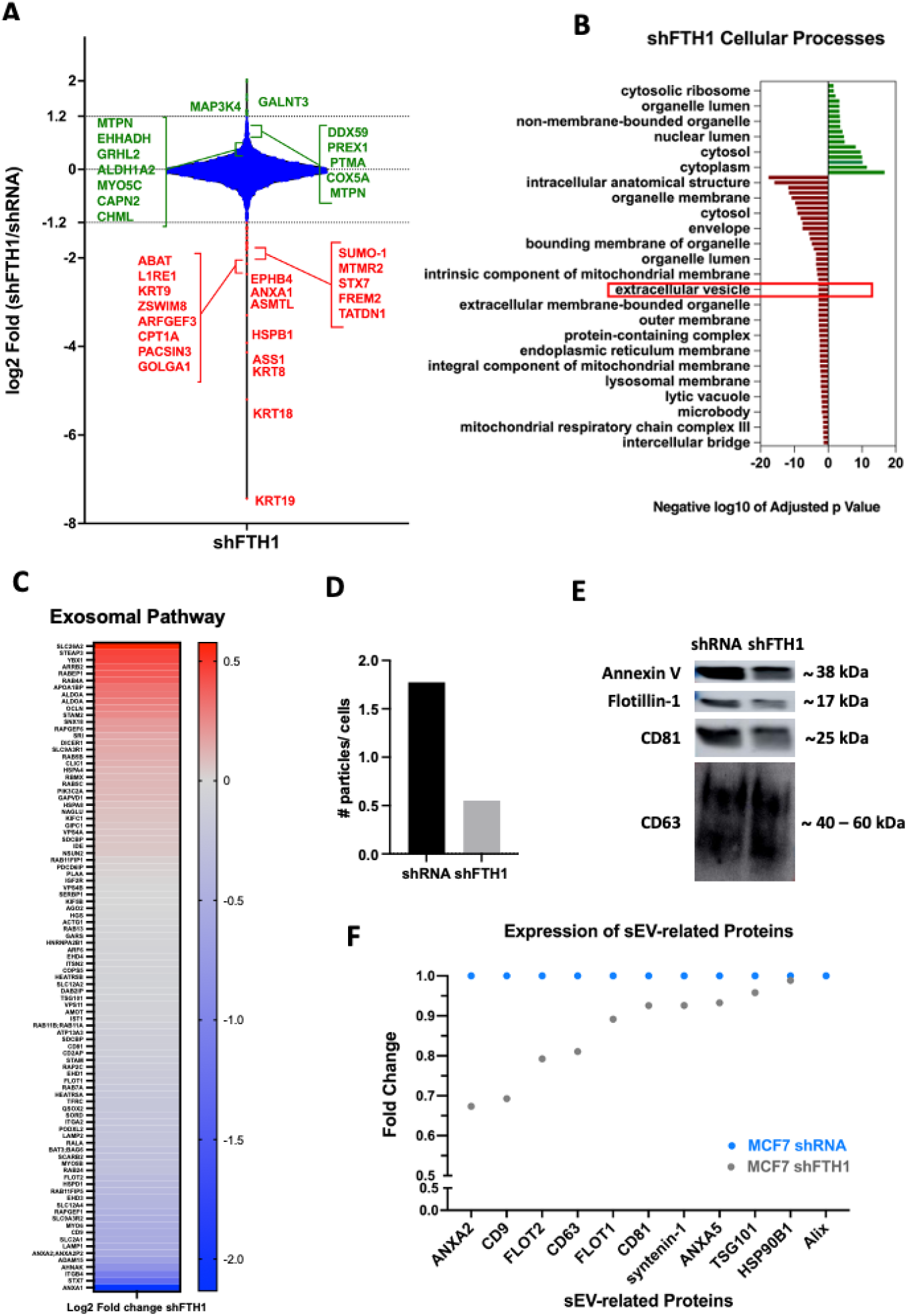
Iron metabolism supports the connection between LD and sEVs. **A**) Violin plot depicting the ratio of Log2 Fold for MCF7 shFTH1/MCF7 shRNA. The proteins for which the expression was highly upregulated (green) or highly downregulated (red) were annotated on the plot **B**) Cellular processes upregulated (green) and downregulated (red) in MCF7 shFTH1 cells. **C**) Heatmap of proteins belonging to the exosomal pathway. Representation of Log2 Fold change values. **D**) Ratio of particle number per cell for the sEV fraction (F2) released by MCF7 shRNA and MCF7 shFTH1 (F2), using NTA. Data are presented as means (n=1). **E**) Western blot for the sEV pellets (100K) obtained by differential ultracentrifugation combined with SEC for MCF7 shRNA and MCF7 shFTH1 cells. The same sample volume (19.5 μL) was loaded onto the 10% acrylamide gel. Annexin V, Flotillin 1, CD81 and CD63 exosomal markers were used. The results presented here are representative of one independent experiment. **F**) Expression of main exosomal markers (Annexin A2 (ANXA2), CD9, flotillin 2 (FLOT2), CD63, flotillin 1 (FLOT1), CD81, Syntenin-1, Annexin A5 (ANXA5), TSG101, HSP90B1 and Alix) is shown for MCF7 shRNA (blue) and MCF7 shFTH1 cells based on proteomic data.

### LD stimulation increases sEV biogenesis

It has been previously reported by our research group and others that X-ray radiation (17, 24) and acidosis (pH 6.5) (25) promote an enrichment in cancer cells with high LD content (LD^High^). Since the LD inhibition led to a decrease of sEV release, we then decided to evaluate the LD-sEV connection in a context of LD stimulation. We therefore studied the effect of pH variation on MCF7 and H460 cell lines. Both cell lines were incubated with neutral pH (7.4) or in acidic (pH 6.5) conditions for 72 hrs. Afterward, the number of LDs per cell was assessed by confocal microscopy. We confirmed a higher number of LDs/cell in acidosis when compared to neutral media for both cell lines (**Fig 4A**). The isolation of sEVs revealed a higher number of particles released per cell (**Fig 4B**) and a higher protein concentration (**Fig S4A**) in low pH conditioned media. In line with these results, the expression of exosomal markers (CD63, CD9, CD81 and hsc-70) on sEVs isolated from acidic condition was more elevated than the neutral one (**Fig 4C and S4B**). The comparison between the two pH settings was carried out using the same sEV sample volume.

**Figure 4.**
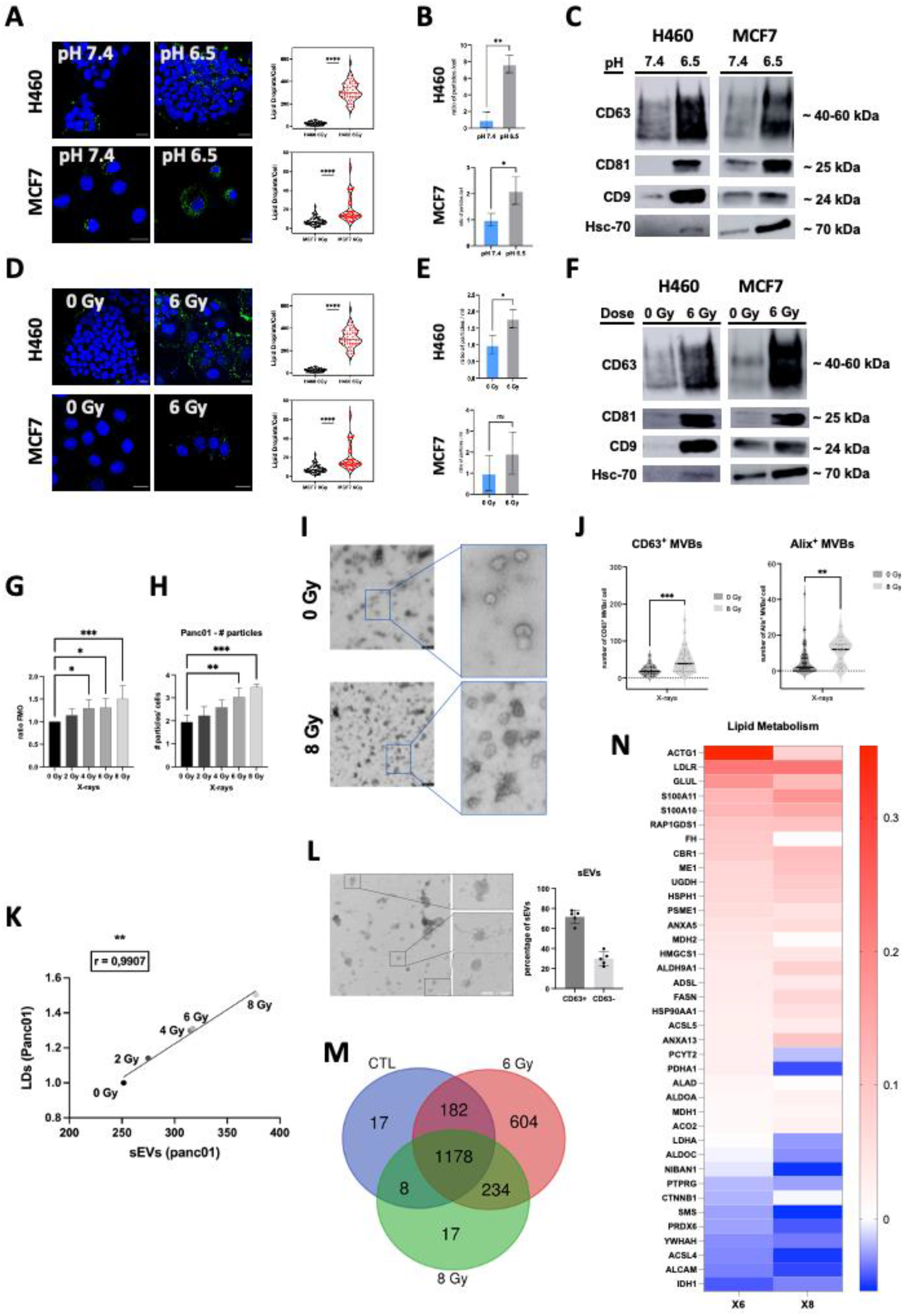
LD stimulation increases sEV biogenesis. **A)** and **D**) Treated (pH 6.5 (**A**) or 6 Gy (**D**)) and untreated (pH 7.4 (**A**) or 0 Gy (**D**)) H460 and MCF7 cells were stained with LD540 (yellow) for LDs and DAPI (blue) for nuclei and imaged at the confocal microscope with a 100X objective (Leica Microsystems; Concord, Ontario, Canada). n=1 (H460 pH 7.4, N=354 cells and pH 6.5, N=166 cells; MCF7 pH 7.4, N=93 cells and pH 6.5, N=33 cells; H460 IR 0 Gy and 6 Gy: N = 29 cells; MCF7 IR 0 Gy and 6 Gy: N = 50 cells). The pinhole was set for a slice thickness of 17.4 μm, with an interval between slices of 0.9 μm. Z-projection of the z-stack acquisitions is shown above. Displayed are the merged images of the LD540 and DAPI staining from one independent experiment (Scale bar, 20 μm). The graph represents the changes in LD content for MCF7 and H460 cell lines. Images were analyzed using ImageJ for mean LDs per cell. Comparisons between groups are shown with corresponding p-values (unpaired Student’s t-test). Error bars represent the means ± SD. **B)** and **E**) Ratio of particle number per cell for the sEV fractions (F2) released by treated (pH 6.5 (**B**) or 6 Gy (**E**)) and untreated (pH 7.4 (**B**) or 0 Gy (**E**)) H460 or MCF7 cells, using NTA. Results from three independent experiments. Data are presented as means ± SD. Comparisons between groups are shown with corresponding p-value (unpaired Student’s t-test). **C) and F**) Western blot for the sEVs pellets (100K) obtained by differential ultracentrifugation combined with SEC for H460 and MCF7. Same sample volume (19.5 μL) was loaded onto the 10% acrylamide gel. The results presented here are representative of three independent experiments. **G**) Panc01 cells, untreated and irradiated with 2, 4, 6 or 8 Gy, were stained with LD540 for LDs and PI for dead cells, and analyzed by flow cytometry. The graph represents the mean fluorescence intensity (MFI) (irradiated/unirradiated ratio). Comparisons between groups are shown with corresponding p-values (ANOVA I, Dunnett’s post-test). Error bars represent the means ± SD. n=3. **H**) Ratio of particle number per cell for sEV fraction (F2) released by Panc01 irradiated with X-rays (0, 2,4,6 or 8 Gy). Results from three independent experiments. Data are presented as means ± SD. Comparisons between groups are shown with corresponding p-value (ANOVA I, Dunnett’s post-test). **I**) High-resolution transmission electron micrograph of sEVs isolated from unirradiated (0 Gy) or irradiated (8 Gy) Panc01 media taken with Zeiss EM 910 at 100 kV. Uranyl acetate negative staining reveals that purified sEVs have a cup-shaped morphology enclosed by a lipid bilayer. The diameter of sEVs is around 90–100 nm. The presented image has a magnification of 16000 x in TEM mode. The size bars on the image represent 250 nm. **J**) Number of CD63+ or ALIX+ MVBs after irradiation (8 Gy) in Panc01 cells transfected CD63-pHLuorin or ALIX-mCherry plasmids (n=1) **K**) Pearson correlation on mean values was run to determine the relationship sEV and LD number. The correlation factor is 0.9907. **L**) Immunogold CD63 staining of Panc01-derived sEV in control condition and quantification of CD63-positive vesicles. The presented images were taken with Zeiss EM 910 at 100 kV and have a magnification of 16000 x in TEM mode. The size bars on the image represent 250 nm. **M**) Venn diagram of sEV proteomics analysis. Comparison of the proteins regulated for X-ray irradiation (6Gy and 8 Gy) with respect to the proteomics analysis of sEVs obtained from unirradiated Panc01 cells. **N**) Heatmap of proteins belonging to the lipid metabolism pathway. Representation of Log2 Fold change values for 6 and 8 Gy X-rays. * ≤ 0.05; ** ≤ 0.01; *** ≤ 0.001 and ****≤ 0.0001.

The same approach was used to study the radiation effect. In our previous work, we showed that cancer cells surviving to 6 Gy X-rays were characterized by an increase of the LD content 72 hrs after irradiation (17, 24). Starting from this premise, we confirmed those data in H460 and MCF7 cells and extended the study to the Panc01 cell line using either confocal imaging or flow cytometry (**Fig 4D, 4G and S4C**). PI was used to make sure PI^+^ cells were not considered in the flow cytometry analysis. However, since the supernatant was changed every 24h and the PI+ cells was very low (2.37%), we estimated that dead cells were washed away at the moment of the analysis (confocal microscopy or flow cytometry). Particle number and analysis of the exosomal marker expression (CD63, CD9, CD81 and hsc-70) demonstrated that irradiation treatment was also able to increase the sEV secretion (**Fig 4 E, 4F** and **S4D**). Interestingly, the cellular LD content increased proportionally to the radiation dose given to the cells (**Fig 4G** and **S4C**), and we observed the same trend for sEVs release (**Fig 4H**). EM also indicated the elevated number of sEVs collected from Panc01 and H460 72 hrs after 8 or 6 Gy X-rays respectively, as compared to the unirradiated conditions (**Fig 4I** and **S4F**). Interestingly, the particle size was similar between the sEVs isolated from irradiated or unirradiated cells (**Fig S4G**). In addition, the exosomal nature of Panc01-derived vesicles was demonstrated by an analysis of CD63^+^ or Alix^+^ multivesicular bodies (MVBs) in unirradiated (0 Gy) or irradiated (8 Gy) pancreatic cancer cells (**Fig 4J**). Moreover, we confirmed a clear correlation between cellular LD content and sEV biogenesis, as represented in **Fig 4K**. Since irradiation induces apoptosis and autophagy, it is important to consider that very small apoptotic bodies (100 – 1000 nm) and autophagic vesicles (40 – 1000 nm) could be co-isolated by differential ultracentrifugation combined with SEC (cut-off 200 nm) within the sEV pool. We therefore characterized the expression of AnnexinV and LC3 on sEVs isolated from Panc01 irradiated cells via western blot (**Fig S4F**) and ELISA (**Fig S4H**), showing an increase expression of those markers. However, an immunogold EM-staining also showed that 71.63% of sEVs were coated by gold-coupled anti-CD63 antibodies in irradiated condition (8 Gy) (**Fig 4L**). Altogether, while we cannot exclude a contamination of our sEVs with small apoptotic and autophagic vesicles after irradiation, we showed that the expression of CD63 on sEVs (western blot), the number of CD63^+^ sEVs (EM) and the number CD63^+^ MVBs (confocal microscopy) were increased after irradiation, meaning that a higher proportion of CD63^+^ vesicles were released.

Finally, to evaluate how irradiation could affect the exosomal cargos,–exosomal proteins were extracted from sEVs either released by X-ray irradiated (6, 8 Gy) Panc01 cells or by their unirradiated counterpart. 431 sEV proteins, analyzed by Mass Spectrometry (**Fig 4M**), were downregulated while 566 proteins had an upregulated expression compared to the unirradiated conditions. Interestingly, a closer look to the lipid metabolism pathway (**Fig 4N**) led us to identify a higher expression of proteins involved in the lipid anabolism in sEVs derived from irradiated Panc01 as compared to the control condition, and especially after 6 Gy. The proteins, whose expression was downregulated, belonged to the lipid catabolism pathway, meaning that irradiation favors lipid biosynthesis while reducing lipolysis, in accordance with the increased LD formation. This also means that radiation, in addition to affect cellular LD content, regulates the lipid-related sEV proteome. This is of high interest since the exosomal lipid proteome and lipid profile modulate the invasiveness of the recipient cells (26–28).

Altogether, these results showed that variation in the tumor microenvironment (e.g., pH), or treatments, such as conventional radiation, can strongly stimulate LD biogenesis and modulate in the same way the interconnected sEV pathway.

### Patient-derived colorectal cancer stem cells modulate their LD content and sEV release under hypoxia

It is known that LDs are considered as a functional marker for cancer stemness (10). Indeed, patient-derived CR-CSCs (**Fig 5A**) with a high LD content exhibited a higher tumorigenic potential (10, 15). Moreover, it was shown that restricted oxygen conditions increased the CSC fraction and promoted the acquisition of a stem-like state (29). Considering this, we decided to study the influence of hypoxia on the LD-sEV interconnection in patient-derived CR-CSCs. By using confocal microscopy, we observed a higher number of LDs/cell when CR-CSCs were cultured in hypoxic conditions as compared to the normoxic state (**Fig 5B**). A parallel NTA analysis showed a higher number of sEVs released by CR-CSCs in hypoxia than in normoxic conditions (**Fig 5C**). The analysis of some exosomal markers also revealed a higher expression of CD9, CD63 and CD81 in hypoxia than in normoxia when the same sEV sample volume was used for western blotting (**Fig 5D**). Overall, we observed a clear correlation between LD content and sEV number with a Pearson’s r coefficient of 0.870 (p < 0.01) (**Fig 5E**).

**Figure 5.**
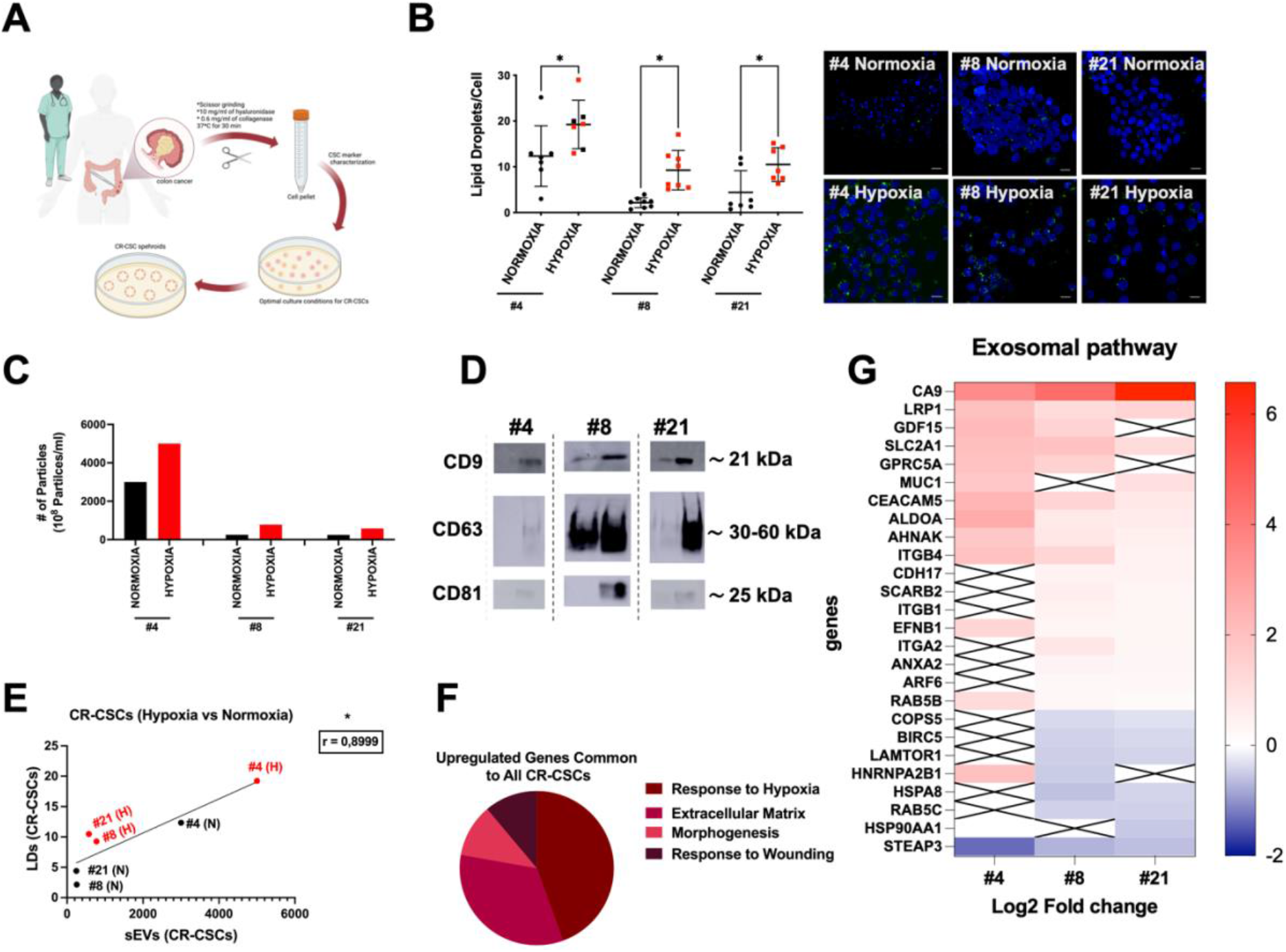
Patient-derived colorectal cancer stem cells modulate their LD content and sEV release under hypoxia. **A)** Schematic representation of CR-CSC isolation and culture**. B**) LD quantification in Colorectal Cancer Stem Cells (CR-CSCs) derived from patients with colorectal cancer. Treated (Hypoxia, N) and untreated (Normoxia, N) CR-CSCs (#4, #21, #8) were stained with BODIPY 493/503 for LDs (green) and DAPI (blue) for nuclei and imaged at the confocal microscope with a 100X objective (Leica Microsystems; Concord, Ontario, Canada). The pinhole was set for a slice thickness of 17.4 μm, with an interval between slices of 0.9 μm. Z-projection of the z-stack acquisitions is shown above. The merged images of the BODIPY and DAPI staining from three independent experiments are displayed (Scale bar, 20 μM). The graph represents the changes in LD content for the different CR-CSCs in hypoxia as compared to normoxia. Images were analyzed using ImageJ for mean LDs per cell. Comparisons between groups are shown with corresponding p-values (ANOVA I, Sidak post-test). Error bars represent the means ± SD. * ≤ 0.05; ** ≤ 0.01; *** ≤ 0.001 and ****≤ 0.0001, n=1. **C**) Ratio of particle number per cell for sEV fraction (F2) treated (Hypoxia, H) and untreated (Normoxia, N) (F2) released by CR-CSCs ((#4, #21, #8). Results from one independent experiment. **D**) Western blot for the sEVs pellets (100K) obtained by differential ultracentrifugation combined with SEC for all CR-CSCs. The sample volume (19.5 μL) was loaded onto the 10% acrylamide gel. The results presented here are representative of one independent experiment. **E**) Pearson correlation on mean values was run to determine the relationship sEV and LD number for CR-CSC when LD content is either high (hypoxia) or low (normoxia). ** ≤ 0.05; ** ≤ 0.01; *** ≤ 0.001 and ****≤ 0.0001. **F**) Diagram of common upregulated pathways in all CR-CSCs culture under hypoxia (Cytoscape: Network specificity 6 genes, 0.4 k score, pValue adjusted <0.05, log2 Fold change >1.2). **G**) Heatmap of proteins belonging to the exosomal pathway. Representation of Log2 Fold change values for the hypoxic condition as compared to the normoxic condition.

Finally, to further evaluate the effect of hypoxia on the lipid metabolism and the exosomal pathway, we collected mRNA from the three different CR-CSCs in normoxia and hypoxia for a full transcriptome analysis. This led us to identify four upregulated pathways under hypoxia using String and Cytoscape: *i)* response to hypoxia; *ii)* extracellular matrix; *iii)* morphogenesis; *iv)* response to wounding (**Fig 5F**).

Downregulated genes belonged to *i*) tRNA pathway and *ii*) positive regulation of double strand break repair via homologous recombination (**Fig S5A**). This analysis also allowed us to confirm that many of the genes involved in the sEV pathway were upregulated (**Fig 5G**) similarly as the sEV number was modulated in the three CR-CSCs (**Fig 5B**). Interestingly, the expression of the genes involved in the lipid metabolic pathways was mainly upregulated under hypoxia, for all CR-CSCs (**Fig S5B**). In general, downregulated lipid metabolism-related genes were associated with lipid catabolism while the upregulated ones were associated with lipid anabolism. As expected, the hypoxia pathway was also upregulated in CR-CSCs cultured under hypoxic as compared to the normoxic conditions (**Fig S5 C).** Altogether, these results showed that the interconnection between LDs and sEVs was also present in patient-derived CR-CSCs cultured in hypoxic conditions.

## Discussion

By modulating cellular LD amount, either through the inhibition of LD metabolism or the stimulation of LD biosynthesis in different cancer cell types, we report for the first time a tight correlation between the intracellular LD numbers and the sEV release. These findings were also validated in patient-derived CR-CSCs showing that hypoxia increased intracellular LDs as well as sEV biogenesis. In addition, multiple omics data confirmed, at the mRNA and protein levels, that LD and sEV pathways were similarly modulated and tightly connected.

It is becoming increasingly clear that LDs are not static organelles involved only in safely storing excessive and dangerous lipids, but they might play a major role as lipid sources for potential membrane-shaped vesicles. While the LD-sEV connection has never been shown so far, hypoxia (29, 31), low pH (6.5) (25, 32, 33), irradiation (17, 34), reactive oxygen species (ROS) (35, 36), high glucose consumption (37–39) and cellular senescence (40, 41), among others, have been shown to stimulate intracellular LD content as well as sEV release by cells. Several studies contributed to elucidate the mechanism behind the increased sEV biogenesis upon those stimulations. For example, cellular senescence and DNA damaging reagents or radiation were shown to stimulate sEV production through the activation of p53, at least partially (42). Intriguingly, p53 is known to activate the expression of several genes involved in endosome regulation, including Rab5B, Caveolin-1, TSAP6 and Champ4C (a subunit of ESCRT-III) (43, 44). In parallel, p53 was also demonstrated to have an impact on the lipid and iron metabolisms (45). Despite its multiple targets, p53 alone is not enough to fully elucidate the link between LDs on one side and the exosome pathway on the other side. Another example is the regulation of sEV release through ATM activation of the autophagic pathway in hypoxia (46). Hypoxia also triggers LD formation through HIF1a stabilization. However, despite its role in sEV biogenesis under hypoxic condition, the stabilization of HIF1a in normoxia was not sufficient to support its role in sEV production (47). Overall, while the link between autophagy and LDs has already been well established and characterized, little is known about how LDs could fuel sEV biogenesis. It is to note that the LD content modulation via acidosis, radiation or hypoxia is not as straightforward as LD inhibition and each of those stimulation cannot be claimed to be processes that only impact LD or sEVs as they have a global cell impact. However, all the experiments presented here, taken altogether, allowed us to establish a strong LD-sEV connection.

With our proteomic analyses, we identified proteins whose expression was modulated according to the LD content. A focus on the proteins involved in the exosomal pathway allowed us to evidence a potential role of Rab18, Rab1a, Rab5c and Rab7a in the interconnection between LDs and sEVs (**Figure 6; Table S1**).

**Figure 6.**
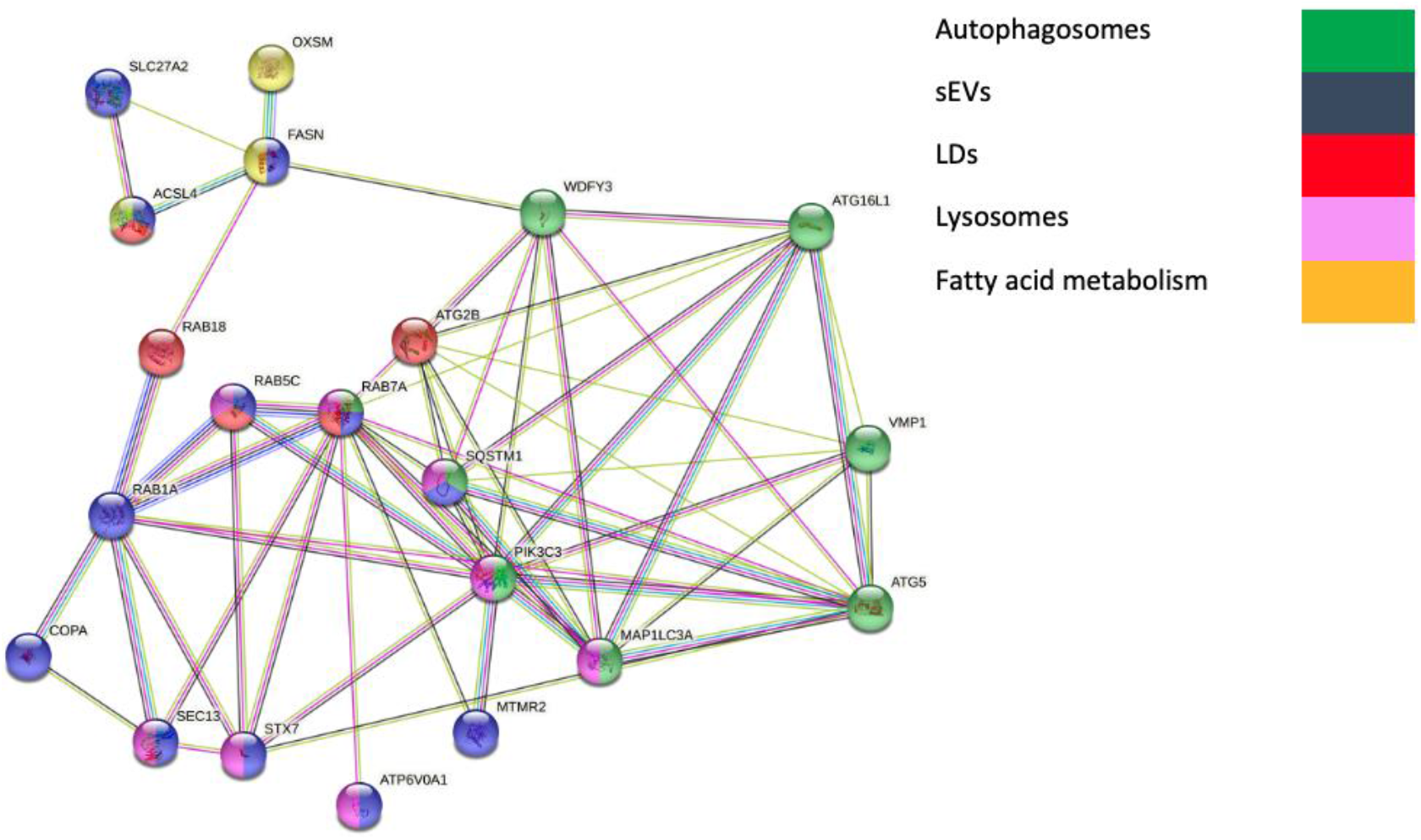
STRING (v.11.5)-based interaction analysis of the proteins identified by mass spectrometry as upregulated in LD^High^ content cells and downregulated in LD^Low^ content cells. A focus on proteins involved in exosome and lipid metabolism allowed to evidence Rab18, Rab5c, Rab7a and Rab1a as key factors in the LD-sEV connection.

In particular, Rab18 knockout was shown to affect the LD growth and maturation, inducing fewer but bigger LDs (48, 49). While Rab18 does not seem to be involved in LD biogenesis, its role in connecting LD catabolism to the autophagic and endosomal pathway is more and more clear (48, 49). Interestingly, Rab18 KO cells showed an increased expression and phosphorylation of ATG2 A/B, ATG9A and ATG16L1, as a compensation to the limited lipid availability (48). In addition, since RAB3GAP1/2 controls the activity and the location of Rab18, its knockout was shown to affect the LD content in the same way as Rab18KO (48, 49). The activity and location of Rab18 on LDs is also controlled by another complex, COPI-TRAPPII (TRAPPC9/TRAPPC10). However, TRAPPII does not seem to play an essential role in the early secretory pathway (50). Finally, Rab18 was found on a sEV subtypes, for which the secretion is mediated by CHMP1A, an ESCRT-III protein (49).

Although the connection of other Rab proteins to the LD and sEV pathways was not extensively investigated, emerging roles of Rab1a, Rab5c and Rab7a in the sEV and LD pathway are recognized. For example, mutations on Rab18, but also Rab5, are known to induce Warburg Syndrome, characterized by the appearance of fewer but bigger LDs (52). Proteomic data published several years ago, also emphasized Rab1a, Rab5b, Rab7a and Rab18 as important players for the connection between LDs and endoplasmic membranes (53). Hypoxia was also shown to increase exosome release via Rab5a (54). In addition, Aromatase inhibitors, through the increased expression of Rab18, Rab5c and rab7a, stimulated the exosome biogenesis (55). However, Rab18, Rab5a, Rab5c and Rab7a were observed on the ectosomes at a higher level than on the sEV/exosomes. On the contrary, Rab1a is more expressed on sEV/exosomes than ectosomes (56). Finally, the investigation of CD63 routes showed its interaction with Rab5 and Rab7 (57). Altogether, while the literature offers insights that support our hypothetical mechanism, further investigations are needed to fully elucidate the LD-sEV connection.

Iron level elevation is associated with ferroptosis, a type of controlled cell death. Interestingly, Ferritin plays a pivotal role in the Fe^2+^ storage (58) and low amount of ferritin drives ferroptosis. Since lipid peroxidation induces ferroptosis (59), cells protect themselves by storing lipids within LDs. Therefore, resistance to ferroptosis is usually associated with LD accumulation. Another way for the cells to deal with an elevated iron level is to promote its export, either through free secretion or via the exosome pathway when associated with ferritin. Indeed, high iron level can trigger CD63 expression via the IRE-IRP pathway and promote the exosomal secretion of ferritin-associated iron (60). In the same context, prominin 2 also favored the exosomal transport of ferritin (61). Interestingly, our results showed that the FTH1 silencing reduced the LD content and the sEV biogenesis, which might increase the iron level and promoting the susceptibility to ferroptosis (58). Since the cells containing the highest LD content are adipocytes, it is also interesting to correlate the sEVs released by those cells. Remarkably adipocytes from obese mice released more exosomes than lean mice (21). In the same context, obesity is associated with a higher risk of carcinogenesis in several organs, including breast, prostate, colon, and liver (62). It also correlates with a faster progression of cancer disease and an increased mortality (27). Interestingly, it was also shown that the fatty acid oxidation (FAO)-related protein content of adipocyte-derived sEVs modified mitochondrial dynamics in recipient melanoma cells, therefore promoting melanoma migration and aggressiveness (63, 64). Similarly, adipocyte-derived exosomes, by transporting neutral lipids, induced an adipose-tissue macrophage phenotype in bone marrow. We showed in our previous publications that LD^High^ cells were more radioresistant than LD^Low^ ones (17, 24). In the present study, we showed that sEVs derived from irradiated cells acquired a stronger lipid biosynthesis profile. Further analyses will help to understand if the lipid-related protein profile of sEVs released upon irradiation also influences the aggressiveness and the metastatic state of targeted cancer cells. At least, it is already known that sEV release after irradiation induced migration and invasiveness in head and neck and breast cancer cells (65, 66).

In conclusion, the possibility to fine-tune sEV biogenesis by targeting LDs could have a vast effect on the amount, the cargos and therefore the properties of sEVs thus potentially having a huge impact in the clinics. Further investigations will also help to shed new light on the mechanistic phenomenon behind the LD-sEV interaction and, by consequence, how to turn these results into a future patient-tailored therapy.

## STAR Methods

### 1. Cell Culture

Different human cancer cell lines, purchased from ATCC, were used in this study. Human colon adenocarcinoma cell lines HT-29 (HTB-38) and LoVo (CCL-229) were cultured in McCoy’s 5A (Modified) Medium, GlutaMAX™ Supplement (1X) (Gibco-Thermo Fischer Scientific, USA; # 36600-021) or Ham’s F-12K (Kaighn’s) Medium Nutrient Mix (1X) (Gibco-Thermo Fischer Scientific, USA; # 21127-022) respectively. Human breast adenocarcinoma cell line MCF7 (HTB-22) was cultured in Dulbecco’s Modified Eagle Medium (DMEM) high glucose (1X) (Gibco-Thermo Fischer Scientific, USA; #11995-065). Human non-small-cell lung carcinoma (NSCLC) cell line NCI-H460 (HTB-177) was cultured in Roswell Park Memorial Institute (RPMI) 1640 Medium (1X) (Gibco-Thermo Fischer Scientific, USA; #22400-089). Human pancreatic epithelioid carcinoma PANC01 (CRL-1469) cell line was cultured in RPMI 1640 Medium (1X) (Gibco-Thermo Fischer Scientific, USA; #22400-089). All media were supplemented with 10% (v/v) heat inactivated fetal bovine serum (FBS) (Gibco-Thermo Fischer Scientific, USA; #10500-064). Cells were maintained in an incubator 5% CO2 atmosphere at 37°C. Cells were split when a confluence of 90% was reached. All cell line were routinely authenticated (Multiplex human Cell Authentication, DKFZ, Germany).

### 2. Isolation of Cancer Stem Cells from Patients

CR-CSCs were isolated from patients affected by colorectal cancer (CRC) who underwent surgical resection, in accordance with ethical policy of the University of Palermo Committee on Human Experimentation. CR-CSC isolation and characterization were carried out as reported elsewhere (67).

Briefly, CRC samples, after being cut in small pieces, were grinded by surgical scissors at 37 °C for 30 min in DMEM medium supplemented with 10 mg/ml of hyaluronidase (Sigma) and 0.6 mg/ml of collagenase (GIBCO). Cell pellets were, subsequently, cultured in a serum-free Ham’s F-12 Nutrient Mix medium (Thermo Fisher Scientific) using ultra-low attachment cell culture flasks (Corning). CR-CSC samples #4, #8 and #21, growing as spheroids, were mechanically and enzymatically disaggregated by Accutase (Thermo Fisher Scientific), when reached 80% of confluency.

Short tandem repeat (STR) analysis using a multiplex PCR assay, including a set of 24 loci (GlobalFilerTM STR kit, Applied Biosystem, USA), was routinely used to authenticate CR-CSCs and compare them to the parental patient tissues.

### 3. Cell Culture and Transfection

Lentiviral transduced MCF7 were stably transduced with a lentiviral DNA containing either an shRNA that targets the 196–210 region of the FTH1 mRNA (sh29432) (MCF-7shFTH1) or a control shRNA without significant homology to known human mRNAs (MCF-7shRNA). MCF-7 shRNA and MCF-7 shFTH1 were cultured in DMEM medium (Thermo Fischer Scientific) supplemented with FBS 10% (Thermo Fischer Scientific), puromycin 1□g/ml (Sigma-Aldrich). Cells were maintained at 37°C in a humidified 5% CO_2_ atmosphere.

### 4. sEV-free FBS

Fetal bovine serum (FBS) (Gibco, Carlsbad, CA, USA) was ultra-centrifuged at 100,000×*g* for 18 hrs at 4 °C. FBS supernatant was then filtered through a 0.22 μm filter (Millipore, US A) and used for sEV-related experiments.

### 5. Treatments (pH, irradiation, hypoxia, inhibitors)

To collect sEVs, H460 (1.8×10^6^), MCF7 (1.0×10^6^), PANC01 (1.5 10^6^), HT29 (2.0×10^6^) and LoVo (3.0×10^6^) cells were seeded in their normal medium (penicillin/streptomycin free) in T75 cm^2^ flasks (Greiner CELLSTAR) 24 hrs prior treatment.

In the case of LD staining, H460 (1.0×10^5^), MCF7 (1.0×10^5^), HT29 (1.0×10^5^) and LoVo (1.0×10^5^) cells were seeded onto 12 pre-autoclaved coverslips (Electron Microscopy Sciences, USA) in a 12-well cell culture plate (Greiner CELLSTAR) and cultured in their normal medium supplemented with 100U/ml penicillin/streptomycin (Thermo Fischer Scientific, USA; #15140122). For X-ray irradiation (6 Gy), 3.5×10^5^ cells were seeded for both H460 and MCF7 cell lines while 1.0×10^5^ cells were seeded in control groups.

#### 5.1. pH Treatment

24 hrs after seeding, H460 or MCF7 cells were divided in two groups: *i*) a control group, for which the medium was replaced with fresh adequate pH 7.4 medium; *ii)* a treated group, cultured with medium for which pH was adjusted to 6.5. The pH of both cell media was adjusted just prior the medium replacement to avoid any kind of pH variation due to oxidation. Treated cells were kept in culture for 72 hrs. Fresh medium was replaced every day for LD experiments.

To avoid the presence of exogenous sEVs in experiments intended to collect cancer cell- derived sEVs, cells were washed twice with Dulbecco’s phosphate buffered saline (DPBS) (Sigma-Aldrich, USA; #8537) and sEV-free FBS media was used (penicillin/streptomycin free).

#### 5.2. Irradiation Treatment

24 hrs after seeding, samples with H460 or MCF7 cells were divided in two groups: i) a control group, unirradiated and ii) a treated group, irradiated with 6 Gy X-rays using a MultiRad 225kV (Faxitron, Germany) irradiator. Treated cells were kept in culture for 72 hrs to select only radioresistant cells at the end of the incubation time. Fresh medium was replaced every day for LD experiments. For PANC01 and H460 cells, 2, 4, 6 or 8 Gy were also used.

As for pH treatment, cells were washed with DPBS, and media were supplemented with sEV-free FBS (penicillin/streptomycin free).

#### 5.3. Hypoxia Culturing Conditions

All experiments in hypoxic conditions were conducted by culturing CR-CSCs in a three-gas incubator (Thermo Fisher) at 37°C with a 2% of Oxygen and with 5% CO2 atmosphere for 72 hrs. LD staining and RNA-seq have been carried at the end of the incubation time keeping all samples in hypoxic conditions.

#### 5.4. Lipid Droplet Inhibition

Two different LD inhibitors were here tested: PF-06424439 (a diacylglycerol acyltransferase 2 (DGAT2) inhibitor; Saint Louis, MO, USA, CN-PZ0233) and Triacsin C (a long-chain fatty acyl CoA synthetase inhibitor) (Cayman Chemical, #10007448).

Both treatments were carried out for 24 hrs with 30 μM of PF-06424439 or 10 μM of Triacsin C. Drug solutions were prepared freshly for every replicates. As for other treatments, cells were washed with DPBS and media was supplemented with sEV-free FBS (penicillin/streptomycin free).

### 6. FACS Sorting

HT29 cells were detached with TrypLE™ Express (Gibco, USA, #12604013) and then centrifuged for 5 min at 300 *g*. Cells were thereafter stained with LD540 for 10 min at 37°C in the dark. Both samples were washed with DPBS three times to remove the excess of the dye and then resuspended in the sorting buffer (PBS Ca/Mg-free, BSA 0.5%, EDTA 2 mM and Hepes 15mM).

Two populations were then sorted based on the LD abundance using a FACSAria Fusion Cell sorter (BD Bioscience).

The 10% LD^High^ (most bright) and 10% LD^Low^ (most dim) cells were collected and, soon after were seeded on a coverslip using a cytospin centrifuge (Thermo Shandon Cytospin3, Marshall Scientific, USA). Cells were then fixed with 4% PFA and an anti-CD63 (NOVUS #NBP2-52225, Germany) was used at a 1/1000 dilution in PBS+BSA 1% for 2 hrs. Thereafter, a donkey anti-mouse IgG (H+L) Alexa Fluor 647 (Thermo Fisher #A-31571, USA), used at 1/2000 dilution in PBS+BSA 1% for 1hr allowed us to stain the MVBs within the cells. Finally, cells were stained with 1mg/mL Hoechst 33342 (Thermo Fisher Scientific, CN-H3570) for 20 min before being processed for the optical imaging acquisition.

### 7. Immunofluorescence and confocal microscopy

#### Lipid Droplet Staining

LD variation among the different treatments was assessed by staining the investigated cell samples with two different dyes, depending on the experiment needs: LD540 and Bodipy 493/503 (Thermo Fisher, CN-D2191). Briefly, cells were seeded onto a coverslip and left in culture the time necessary for the experiment endpoints (72 hrs for irradiation, hypoxia and LD inhibition, while only 24 hrs for pH). When ready, cells were washed with DPBS, fixed with 4% PFA and then stained with 0.1 mg/ml LD540 or 2 mM Bodipy, both in DPBS. The volumes of the staining solutions were kept constants for all the analyzed cell samples. Nuclei were stained with 1mg/mL Hoechst 33342 (Thermo Fisher Scientific, CN-H3570).

#### CD63 and Alix plasmid transfection for confocal microscopy

Plasmids mCherry-hAlix (plasmid#21504) and pCMV-Sport6-CD63-pHluorin (plasmid #130902) were purchased from Addgene. Cells were plated at a density of 7.5×10^4^ onto glass coverslips in twelve-well plates and allowed to grow in the incubator for 24h. Then the cells were irradiated (8Gy) and were immediately transfected with the plasmids encoding CD63 or Alix, using FuGENE HD reagent (Promega, E2311, USA) with a FuGENE HD:DNA ratio of 4:1. After 48h post transfection the cells cells were washed with DPBS, fixed with 4% PFA for 10 minutes and then stained with 1 mg/ml Hoechst 33342. The images were taken exactly as mentioned in the above paragraph.

#### Confocal microscopy

Whole z-stacks images for the stained cells were taken by using a Zeiss LSM710 or Leica SP5 confocal microscope systems equipped with a 40x (lipid droplets) or 63x (multivesicular bodies, MVBs) oil immersion i-Plan Apochromat (numerical aperture 1.40) objectives. LD540 and Bodipy 493/503 were visualized using the 488 nm laser excitation and a 505-530 nm band-pass filter.

### 8. Lipid Droplet Staining for Flow Cytometry Analysis

Briefly, 1.5×10^6^ cells were seeded into T75 cm^2^ flasks (Greiner CELLSTAR) 24 hrs prior irradiation (2, 4, 6, 8 and 10 Gy) and left in culture for 72 hrs after irradiation. Cells were detached with TrypLE™ Express (Gibco, USA, #12604013) and then centrifuged for 5 minutes at 300x*g*. Cells were thereafter stained with 0.1 mg/ml LD540 for 10 min at 37°C in the dark. Samples were washed with DPBS three times in order to remove the excess of the dye and then resuspended in the sorting buffer (PBS Ca/Mg-free, BSA 0.5%, EDTA 2 mM and Hepes 15mM). PI (Sigma-Aldrich, #P4864, Germany) was used to stain dead cells. Finally, the samples were analyzed using a FACS Canto II (BD Biosciences, USA).

### 9. Differential Centrifugation and sEV Isolation by Size Exclusion Chromatography

Collected supernatants were supplemented with 1 mM Phenylmethylsulfonyl Fluoride (PMSF - Serva, Germany; # 32395) and 100U/ml penicillin/streptomycin (Thermo Fischer Scientific, USA; #15140122) before being centrifuged at 300×*g* for 10 min at 4°C in a swing-out centrifuge to remove cellular debris. Resulting 2,000×*g* supernatants were transferred into ultracentrifugation tubes (Thin-wall, Polyallomer 38.5 ml tubes, Beckman Coulter, USA; #326823) and centrifuged at 100,000×*g* for 2 hrs at 4°C using a Beckman L8-55MV ultracentrifuge (Beckman Coulter GmbH, Krefeld, Germany) with a SW27 Swinging-Bucket Rotor. Resulting 100,000×*g* pellets were resuspended in 200 μL of 0.22-μm-filtered PBS. Size exclusion chromatography was then used to separate the sEVs from the contaminants (e.g., proteins), as previously reported (22).

Briefly, single qEV 35 nm columns (Izon, Christchurch, New Zealand) were allowed to reach room temperature for 30 min. The resuspended pellet fraction (200 μL) was added onto the column. As soon as the sample volume was taken up by the column, 0.22 μm-filtered PBS was added to the top of the column tube. The following fractions were collected: F0 (800 μL = void volume of the column) and F1 to F7 (200 μL each), according to the manufacturer’s instructions.

### 10. Protein Extraction and Quantification (Cells and sEVs)

#### Bicinchonic Acid

Protein concentration of cell samples was assessed employing Pierce™ BCA Protein Assay Kit (Thermo Fisher Scientific Inc., Waltham, MA, USA). Cells were lysed in 300 μL of 1× RIPA buffer (Abcam, Cambridge, UK) supplemented with Halt™ Protease Inhibitor Cocktail, EDTA-free (100X) (Thermo Fisher, USA; #78425) and Halt™ Phosphatase Inhibitor Cocktail, (100X) (Thermo Fischer, USA, #78428). Samples were incubated for 20 min on ice and then centrifuged at 17,000×*g* for 20 min at 4 °C. Resulting supernatants were subjected to BCA assay according to the manufacturer’s instructions. Absorbance was assessed at 562 nm with the use of a plate reader.

#### Qubit

To determine the protein concentration of the isolated sEV samples, Qubit Protein Assay Kit (Life Technologies, USA) was used. SDS (Thermo Fisher Scientific, DE) was used to extract proteins. Briefly, 0.8 μL SDS 2% and 7.2 μL sEV sample were added in labeled Qubit assay tubes and vortexed for 30 sec. The resulting samples were then processed according to the manufacturer’s instructions. For the standards (Qubit™ protein standard #1, #2, #3), 0.8 μL SDS 2% and 10 μL standards were added to the corresponding labelled Qubit tubes.

### 11. Nanoparticle Tracking Analysis (NTA)

Particle quantification of sEV samples was performed via NTA using NanoSight LM10 equipped with a 405 nm laser (Malvern Instruments, Malvern, UK). For the NTA analysis, samples were diluted 1:250 in 0.22 μm-filtered PBS. Camera level and detection threshold were set up at 13 and 1.8, respectively. The absence of background was verified using 0.2 μm-filtered PBS. For each sample, five videos of 40 sec each were recorded and analyzed using the NTA 3.0 software version (Malvern Instruments, Malvern, UK).

### 12. Immunoblotting

sEVs were lysed in RIPA Lysis and Extraction Buffer 10X (Cell Signaling Technology, USA #98010) for 20 min on ice. Per lane, 19.5 μl of protein samples were loaded onto 10% polyacrylamide gels. Following SDS-PAGE and protein transfer, membranes were blocked in 5% bovine serum albumin in PBS-Tween 0.1%, and primary antibodies against CD63 (1:1,000, Novus # NBP2-42225), CD81 (1:1000, ProSci Inc., San Diego, CA, USA, #5195), CD9 (1:1000, Cell Signaling Technology, Danvers, MA, USA, #13174) and hsc-70 (1:1000 Santa Cruz # sc-7298) were used to detect sEV markers.

Calnexin (1:500, GeneScript, Piscataway, NJ, USA, #A0124040), Cytchrome C (1:750, GeneScript, Piscataway, NJ, USA, #A0150740), GM130 (1:1000, Cell Signaling Technology, Danvers, MA, USA, #12480) and Enolase 1 (ENO-1) (1:1000, Abgent, San Diego, CA, #AP6526c) were used in indicated dilutions in 5% BSA in PBS-Tween 0.1% when cell proteins were compared to sEV ones, in order to exclude possible contaminants in sEV fractions.

Either HRP-linked Goat anti-Rabbit (Cell Signaling, USA; #7074), HRP-linked Goat anti-Mouse (Cell Signaling, USA; #7076) or HRP-linked Goat anti-Mouse (Thermo Fisher Scientific, USA; #631462) were used as secondary antibodies. Signals were visualized after secondary antibody hybridization by chemiluminescence detection reagent (Bio-Rad Lab, Hercules, CA, USA, #1705061) with Amersham Imager 680 (GE Healthcare, USA).

### 13. Electron Microscopy (EM)

For negatives staining EM, sEV fractions (F2) were adsorbed onto pure carbon-coated EM-grids for 5 min, washed in aqua bidest and negatively stained with 1% aqueous uranyl acetate. For immuno-EM, sEV fractions were adsorbed on formvar-carbon-coated EM-grids. The incubation with primary antibody (anti-CD63, 1:1000, BD Pharmingen, USA, #556019) was performed after buffer wash and incubation with blocking agent (Aurion, Wageningen, The Netherlands). Protein A-Au was used as reporter (CMC, UMC Utrecht, The Netherlands, size of Au-grains 10nm). Micrographs were taken with a Zeiss EM 910 or EM 912 at 80 kV (Carl Zeiss, Oberkochen, Germany) using a slow scan CCD camera (TRS, Moorenweis, Germany).

### 14. RNA Sequencing Analysis

Total RNA was extracted by RNeasy Mini Kit (Qiagen) and mRNA libraries were prepared using TruSeq® Stranded mRNA Library Prep. Next-Generation Sequencing (NGS) technology (RNA-seq) was used to identify some vital biological processes and pathways involved in fatty acid modulation on CSCs cultured in Hypoxia and Normoxia. Illumina HiSeq 4000 and NovaSeq 6000 were used to perform transcriptome sequencing. The reads were aligned to GRCh38/hg38 of the human genome using STAR version 2.6.1d. Alignments were validated using a combination of FastQC version 0.11.8, SAMtools version 1.9, and MultiQC version 1.7 (68, 69). Transcript abundance estimation was further performed using Salmon version 0.14.1 followed by importing them at the gene level with tximport version 1.14.0 (70, 71). Subsequently, expression analysis at the gene level was conducted with DESeq2 version 1.26.0 (72). Targeted gene analysis of commonly known genes and MORPHEUS Versatile matrix visualization and analysis software were used to visualize the datasets as heat maps (Morpheus, https://software.broadinstitute.org/morpheus).

### 15. Proteomic Analyses

#### Cells

MCF7-shRNA and MCF7-shFTH1 cells were washed twice and then scraped into 2ml of cold PBS. Cells were then centrifuged at 300x*g* for 5 min. Each pellet was incubated with 1mL of 1X Ripa Buffer (Cell Signaling) additioned with HaltTM Protease Inhibitor Single-Use Cocktail, (Thermo Fisher Scientific) and HaltTM Phosphatase Inhibitor Single Use Cocktail (Thermo Fisher Scientific), both diluted 1:100 for 10 min on ice. Lysates were then sonicated (40% amplitude, 10 s/cycle; 3 cycle; 4°C) and incubated for 15 min on ice. 100ml of Benzonase 2,75 U/ml (Millipore-Novagen) was added to lysates, incubated in ice for 10 min and then centrifuged at 2,500x*g* for 30 min at 4°C. The supernatants were collected. Protein concentration was measured by BCA Protein assay kit (Thermo Fisher Scientific) at 562 nm.

#### sEVs

The protein quantification was performed with Qubit assay as described in section 10 (protein quantification).

#### Sample Processing

Samples were thawed and extensively vortexed before proceeding. Subsequently, for each sample, 10 μg protein were processed in a 1 μg/ 3 μL concentration in 1 % SDS and 100 mM ammonium bicarbonate (ABC, Sigma-Aldrich). In brief, 10 mM TCEP, 40 mM chloroacetamide (CAA), 100 mM ABC, and 1x protease inhibitor cocktail (PIC, cOmplete, Sigma-Aldrich) were added to each sample, followed by incubation at 95°C for 5 minutes. Protein binding to Sera-Mag Speed Beads (Fisher Scientific, Germany) was induced by increasing the buffer composition to 50% acetonitrile (ACN, Pierce – Thermo Scientific). The bead stock was prepared as follows: 20 μL of Sera-Mag Speed Beads A and 20 μL of Sera-Mag Speed Beads B were combined and rinsed with 1x 160 μL ddH2O, 2x with 200 μL ddH2O, and re-suspended in 20 μL ddH2O for a final working stock of which 2 μL were added per sample. The autoSP3 protein cleanup was performed with 2x ethanol (EtOH, VWR International GmbH, Germany) and 2x ACN washes. Reduced and alkylated proteins were digested on-beads and overnight at 37°C in a lid-heated PCR cycler (CHB-T2-D ThermoQ, Hangzhou BIOER Technologies, China) in 100 mM ABC with sequencing-grade modified trypsin (Promega, USA). Upon overnight protein digestion, each sample was acidified to a final concentration of 1% trifluoroacetic acid (TFA, Biosolve Chimie). MS injection-ready samples were stored at −20°C.

#### Data Acquisition and Processing

For the data acquisition a timsTOF Pro mass spectrometer (Bruker Daltonics) was equipped with an Easy nLC 1200 system (Thermo). An equivalent of 200 ng protein per sample was injected using the following method: peptides were separated using the Easy nLC 1200 system fitted with an analytical column (Aurora Series Emitter Column with CSI fitting, C18, 1.6 μm, 75 μm x 25 cm) (Ion Optics). The outlet of the analytical column with a captive spray fitting was directly coupled to a timsTOF Pro (Bruker) mass spectrometer using a captive spray source. Solvent A was ddH2O (Biosolve Chimie), 0.1% (v/v) FA (Biosolve Chimie), and solvent B was 100% ACN in dH2O, 0.1% (v/v) FA. The samples were loaded at a constant pressure of 800 bar. Peptides were eluted via the analytical column at a constant flow of 0.4 μL per minute at 50°C. During the elution, the percentage of solvent B was increased in a linear fashion from 2 to 17% in 22.5 minutes, then from 17 to 25% in 11.25 minutes, then from 25 to 37% in 3.75 minutes, and from 37% to 80% in a further 3.75 minutes. Finally, the gradient was finished with 3.75 minutes at 80% solvent B. Peptides were introduced into the mass spectrometer via the standard Bruker captive spray source at default settings. The glass capillary was operated at 1600 V and 3 L/minute dry gas at 180°C. Full scan MS spectra with mass range m/z 100 to 1700 and a 1/k0 range from 0.85 to 1.3 V*s/cm2 with 100 ms ramp time were acquired with a rolling average switched on (10x). The duty cycle was locked at 100%, the ion polarity was set to positive, and the TIMS mode was enabled. The active exclusion window was set to 0.015 m/z, 1/k0 0.015 V*s/ cm2. The isolation width was set to mass 700-800 m/z, width 2 – 3 m/z and the collision energy to 1/k0 0.85-1.3 V*s/ cm2, energy 27-45 eV.

The resulting raw files were searched using MaxQuant version 2.0.3.0 using the default settings unless otherwise stated. Label-free quantification (LFQ) and intensity-based absolute quantification (iBAQ) were applied using the default settings. Matching between runs was enabled. The resulting proteinGroups and peptide tables were further analyzed using matrixQCvis and R.

Protein analysis of commonly known proteins was performed using STRING (https://string-db.org; v.11.5) and cytoscape (v. 3.9.1).

### 16. Statistical Analysis

#### Image analysis

Twelve-bit z-stack images were acquired and post-processed for the LD quantification as reported elsewhere (17). Briefly, the background was subtracted from all images using ImageJ’s Rolling ball radius tool. After that, all images were processed with Gaussian filter, thresholded and segmented with Find Maxima tool. At this point, processed images were analyzed with Analyze Particle tools. The whole image processing was set up automatically thanks to the in-house developed FiJi macro generously provided by Dr. Damir Krunic. Statistical analysis was performed by Student’s t-test with unequal variances. Only p-values below 0.05 were considered statistically significant between two groups.

#### sEVs

Results of the functional analysis were analyzed for statistical significance with GraphPad PRISM 8.0 software (GraphPad Software, San Diego, CA, USA), using unpaired t-test or one-way analysis of variance (ANOVA), followed by Tukey’s multiple comparisons. The differences between means were considered significant if *p* ≤ 0.05. The results are expressed as the means ± standard deviation.

## Acknowledgements

We gratefully acknowledge the imaging and FACS Facilities at the DKFZ and the CoreLab Genomic Facility at KAUST for their prompt and precious support. We also are grateful to the Dr. Sebastian Dieter’s group for the continuous access to the ultracentrifuge.

Carlo Liberale and Joao Seco acknowledge funding from King Abdullah University of Science and Technology, Grant Award Number: OSR-CRG2018-3747.

Luca Tirinato has received funding from AIRC and from the European Union’s Horizon 2020 Research and Innovation Programme under the Marie Sklodowska-Curie grant agreement n. 800924.

Jeannette Jansen was supported by grants of the German-Israeli Helmholtz Research School in Cancer Biology – Cancer Transitional and Research Exchange Program (Cancer-TRAX).

Daniel Garcia-Calderon was funded by the Graduate School Scholarship Programme, 2019 from the DAAD.

## Author Contributions

Geraldine C. Genard: Conceptualization, Methodology, Data Curation, Validation, Formal Analysis, Investigation, Writing Original Draft. Luca Tirinato: Conceptualization, Methodology, Data Curation, Validation, Formal Analysis, Investigation, Writing Original Draft and Funding Acquisition. Francesca Pagliari, Jessica Da Silva, Alessandro Giammona, Fatema Alquraish, Marie Bordas and Maria Grazia Marafioti: Methodology and Investigation. Simone Di Franco, Jeanette Jansen, Daniel Garcia, Rachel Hanley, Clelia Nisticò, Yoshinori Fukusawa Torsten Müller and Jeroen Krijgsveld: Data Curation. Matilde Todaro, Francesco Saverio Costanzo, Giorgio Stassi and Kendra K Mass: Supervision, Investigation and Funding Acquisition. Michelle Nessling and Karsten Richter: TEM Data Curation and Validation. Carlo Liberale: Project Supervision, Funding Acquisition, Data Curation and Draft Revision. Joao Seco: Conceptualization, Methodology, Funding Acquisition, Project Supervision and Draft Revision.

