## Supplemental Material for "Lipid Droplets Fuel Small Extracellular Vesicle Biogenesis"

**Supportive Information**

#### Supportive Figure 1

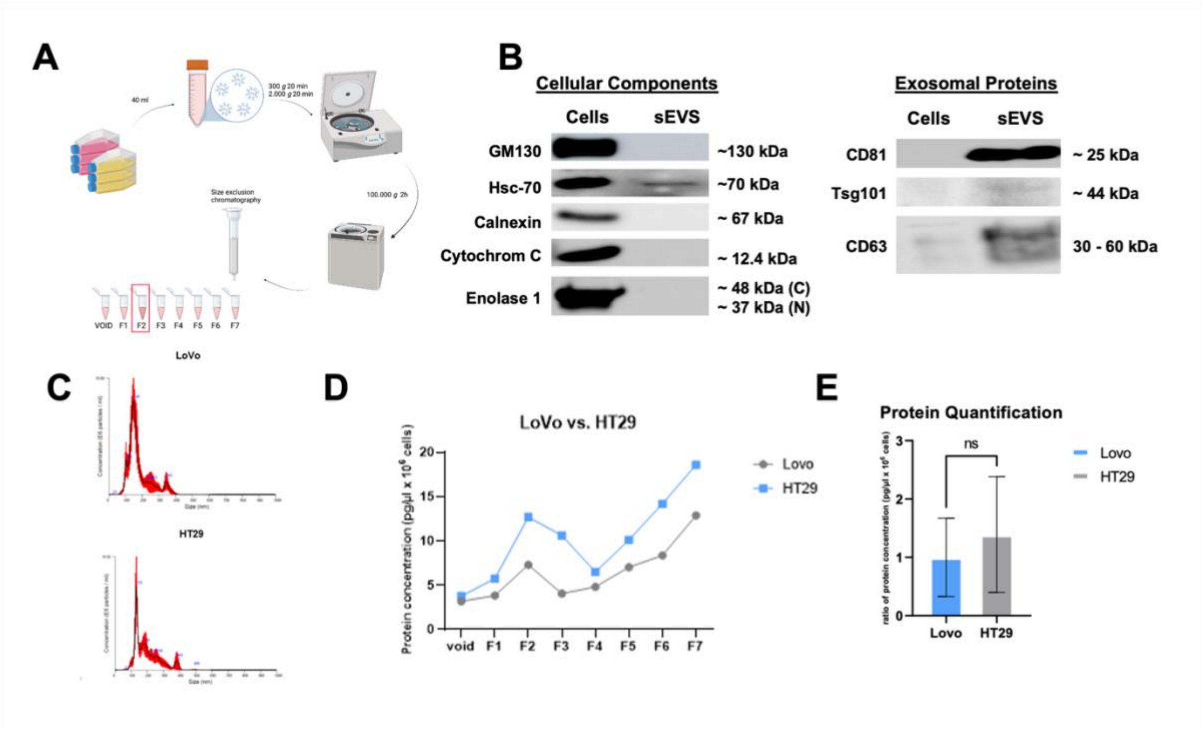


**Figure S1**. **Analysis of LDs content and sEV release in two colorectal cancer cell lines, LoVo and HT29. A**) Summary of sEV isolation by ultracentrifugation combined to SEC (Created with Biorender). **B**) Western blot evaluating the protein expression in cells and sEVs pellets. The same amount of protein (3.5 µg) was loaded onto the 10% acrylamide gel. Cellular components such as Golgi (GM130), Endoplasmic Reticulum (Calnexin-C), mitochondria (Cytochrome C) and other cytoplasmic proteins (Hsc-70 and enolase-1) were observed in the cell samples while sEV proteins, such as CD81, Tsg101 and CD63, were enriched in the sEV samples. **C**) Representative particle size and concentration distribution profiles for isolated IZON fraction 2. Graph results represent a mean of five recorded videos of 30 sec for each sample for LoVo (upper graph) and HT29 (lower graph). **D**) Comparative isolation of size-exclusion column fractions for LoVo (grey) and HT29 (blue). SEC-serial fractions (F0 = void volume, 800µl; F1, F2, F3, F4, F5, F6, F7 (volume 200 µL)) and subsequent fluorometric protein quantification values. Protein concentration for each fraction is shown in a picogram per microliter per 1 million cells (pg/µL/$10^6$ cells). **E**) Protein quantification of F2 fraction for LoVo and HT29 (in pg/µL/$10^6$cells).

### Supportive Figure 2

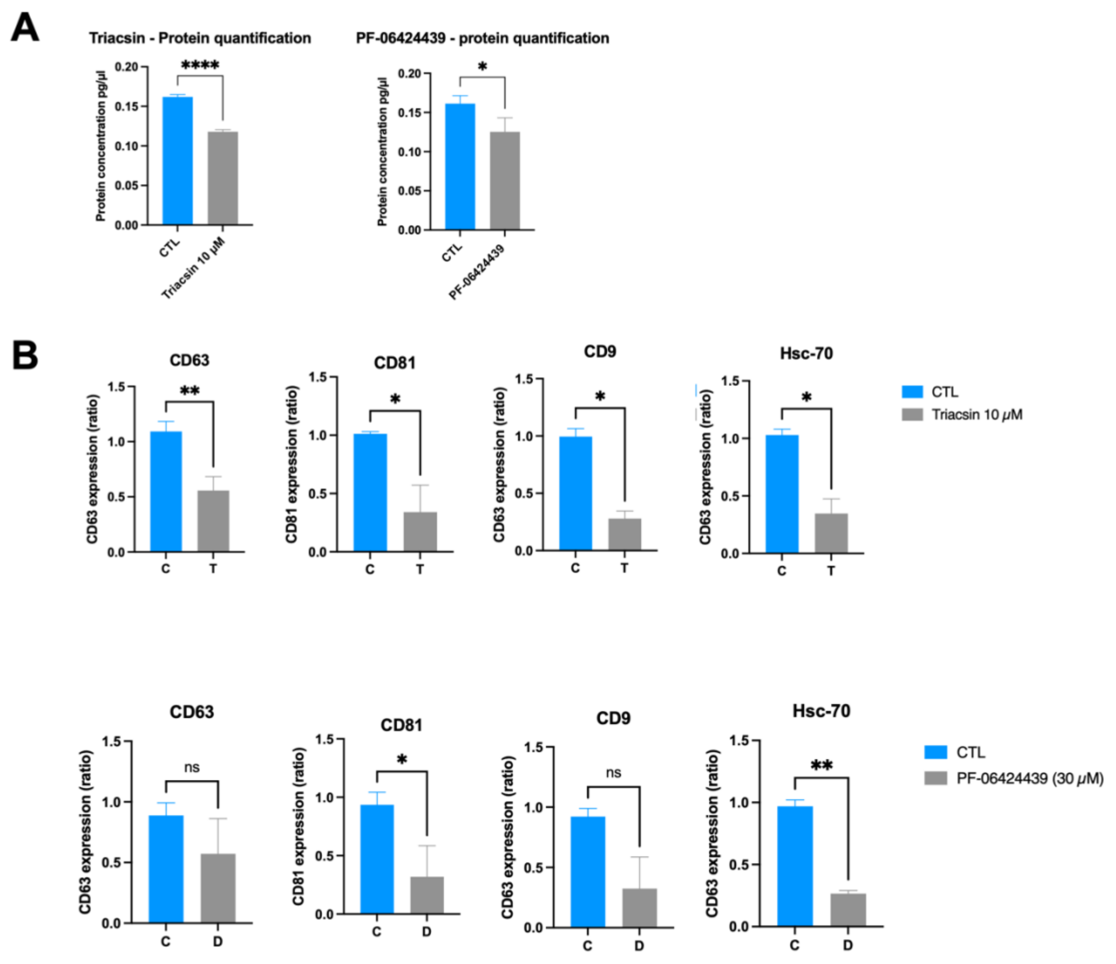


**Figure S2**. **Inhibition of LD metabolism reduces sEV release A**) Protein quantification of F2 fraction for HT29 control or treated, either with Triacsin C or PF-06424439 (in pg/µL/$10^6$cells). **B**) Western blot for the sEV pellets (100K) obtained by differential ultracentrifugation combined with SEC for HT29 cells control or treated, either with Triacsin C 10 µM or PF-06424439 30 µM. The same sample volume (19.5 µL) was loaded onto the 10% acrylamide gel. The graph represents the average of three independent experiments. The intensity of the bands corresponding to treated HT29-derived proteins was normalized by the intensity of the untreated HT29-derived protein band. Unpaired students t-test was performed. Error bars represent the means ± SD. * ≤ 0.05; ** ≤ 0.01.

#### Supportive Figure 3

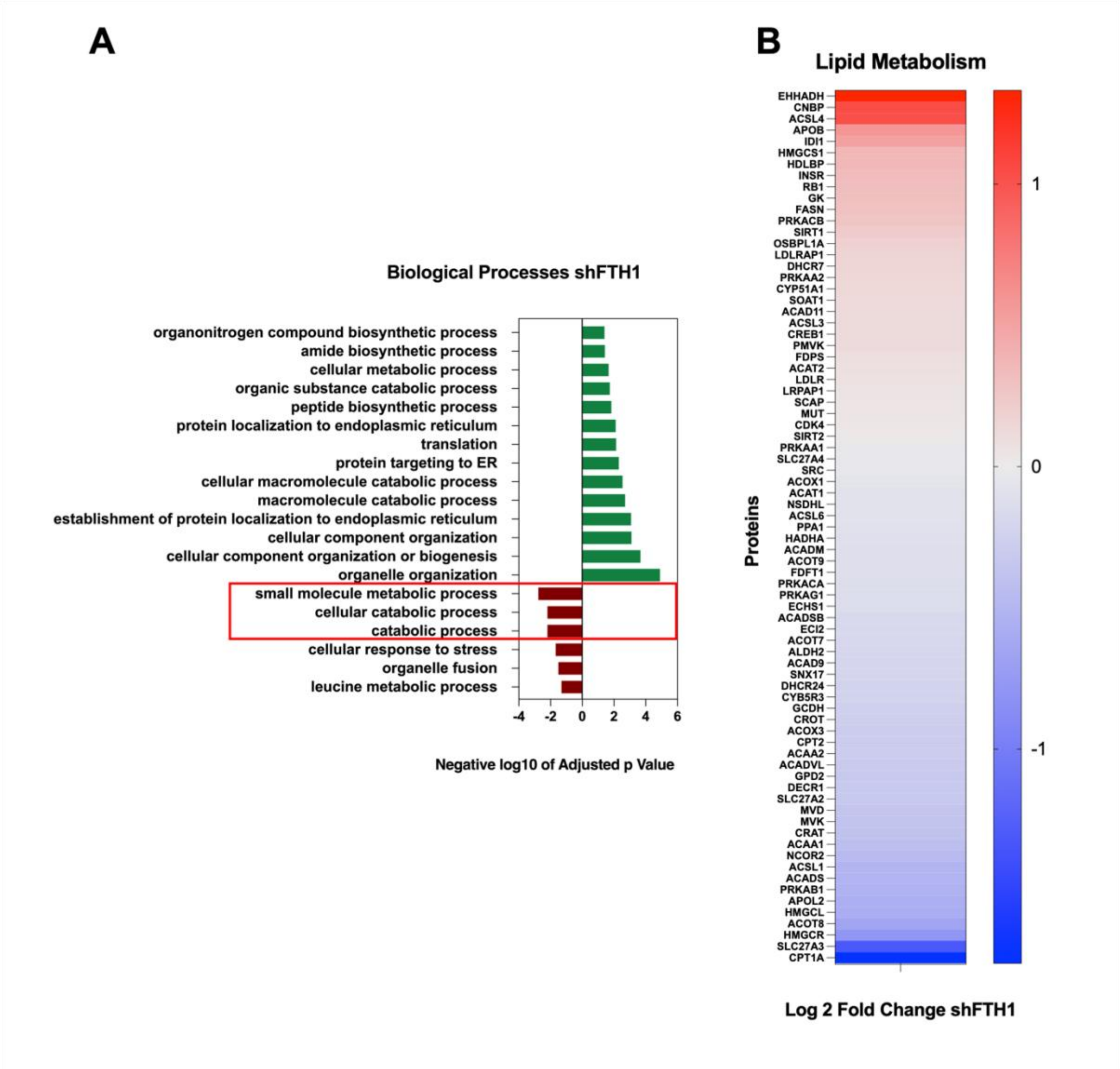


**Figure S3**. **Iron metabolism inhibition supports the connection between LD and sEVs. A**) Biological processes upregulated (green) and downregulated (red) in MCF7 shFTH1. **B**) Heatmap of proteins belonging to the lipid metabolism pathway. Representation of Log2 Fold change values.

#### Supportive Figure 4

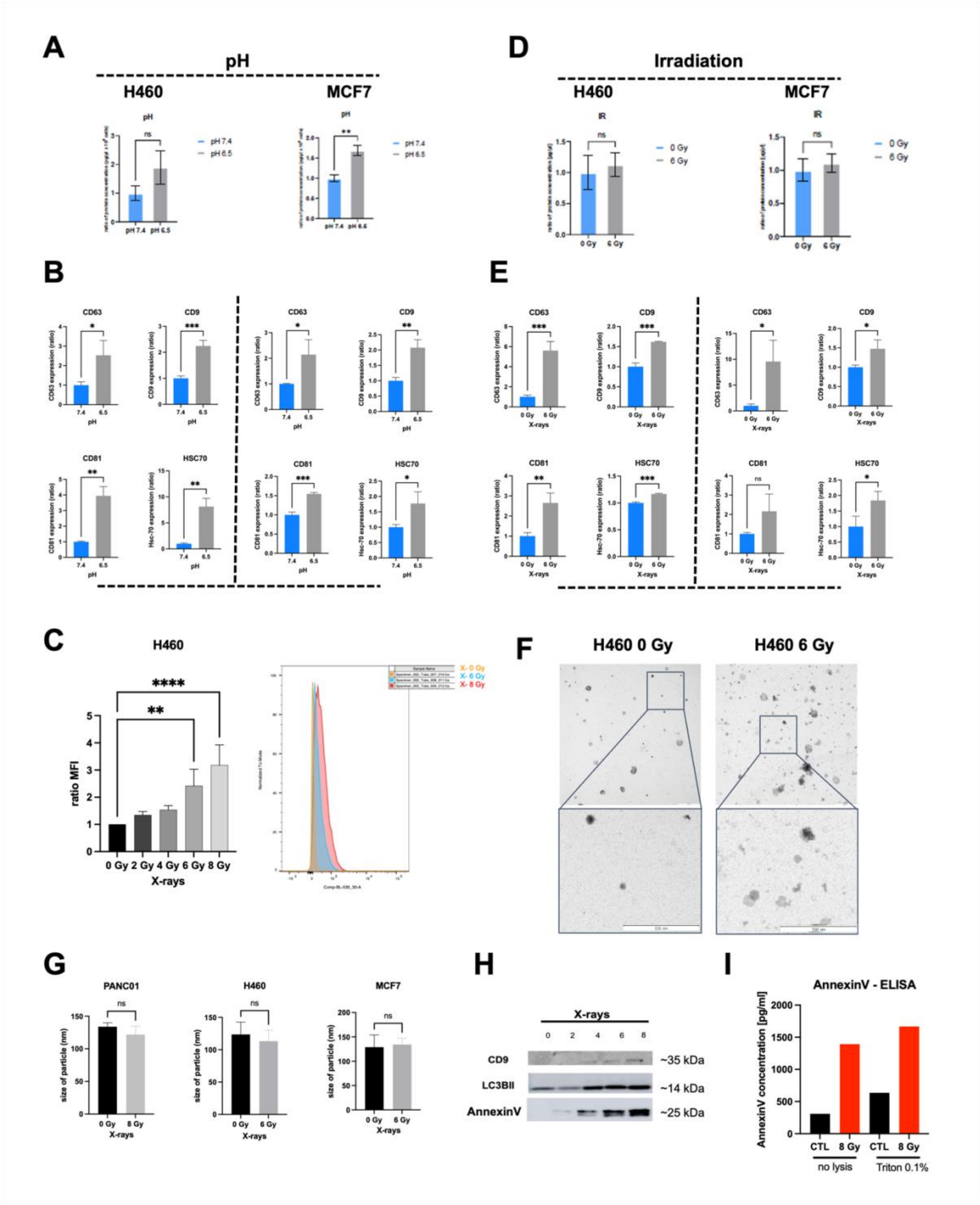


**Figure S4**. **LD stimulation increases sEV biogenesis. A**) Protein quantification of sEV fractions (F2) for H460 (left) and MCF7 (right) treated with neutral (7.4) or acidic (6.5) pH (in pg/µL/$10^6$cells). **B**) Quantification of western blot from three independent experiments. The intensity of the bands, corresponding to the exosomal markers (CD63, CD81, CD9 and Hsc-70) from MCF7 or H460 cultured in acidic (6.5) pH, was normalized by the intensity of the proteins band in control condition (pH 7.4). Unpaired students t-test was performed. Error bars represent the means ± SD. **C**) H460 cells, irradiated with 2, 4, 6 or 8 Gy, were stained with LD540 for LDs and compared with the untreated ones. PI staining was used for discriminating between live and dead cells (less than 3%). Stained cells were analyzed by flow cytometry. The graph represents the mean fluorescence intensity (MFI) (irradiated/unirradiated ratio) for H460 cells. Comparisons between groups are shown with corresponding p-values (ANOVA I, Dunnett's post-test). Error bars represent the means ± SD, n=3. **D**) Protein quantification of sEV fractions (F2) for H460 (left) and MCF7 (right) irradiated (6 Gy) or not (0 Gy) (in pg/µl/$10^6$cells). **E**) Quantification of western blot from three independent experiments. The intensity of the bands, corresponding to the exosomal markers (CD63, CD81, CD9 and hsc-70) from MCF7 or H460 irradiated (6 Gy), was normalized by the intensity of the proteins band in the control condition (0 Gy). Unpaired students t-test was performed. Error bars represent the means ± SD **F**) High-resolution transmission electron micrograph of sEVs isolated from unirradiated (0 Gy) or irradiated (6 Gy) H460 media taken with Zeiss EM 910 at 100 kV. Uranyl acetate negative staining reveals that purified sEVs have a cup-shaped morphology enclosed by a lipid bilayer. The presented image has a magnification of 16000 x in TEM mode. The size bars on the image represent 500 nm. **G**) Representative particle size for isolated IZON fraction 2 of irradiated (6 Gy or 8 Gy) or unirradiated (0 Gy) Panc01 (left), H460 (middle) and MCF7 (right) cells. Error bars represent the means ± SD, n=3 **H**) Western blot for the sEVs pellets (100K) obtained by differential ultracentrifugation combined with SEC for sEV sample volume (3µg) was loaded onto the 10% acrylamide gel. The sEVs were collected from the supernatant of Panc01 cells irradiated with different X-ray doses. The expression of sEV (CD9, AnnexinV), autophagic (LC3-II) and apoptotic (AnnexinV) markers was analysed then analysed. **I**) AnnexinV ELISA with sEV fraction of sEV lysate determining the AnnexinV concentration (pg/ml). * ≤ 0.05; ** ≤ 0.01; *** ≤ 0.001 and ****≤ 0.0001.

**Supportive Figure 5**

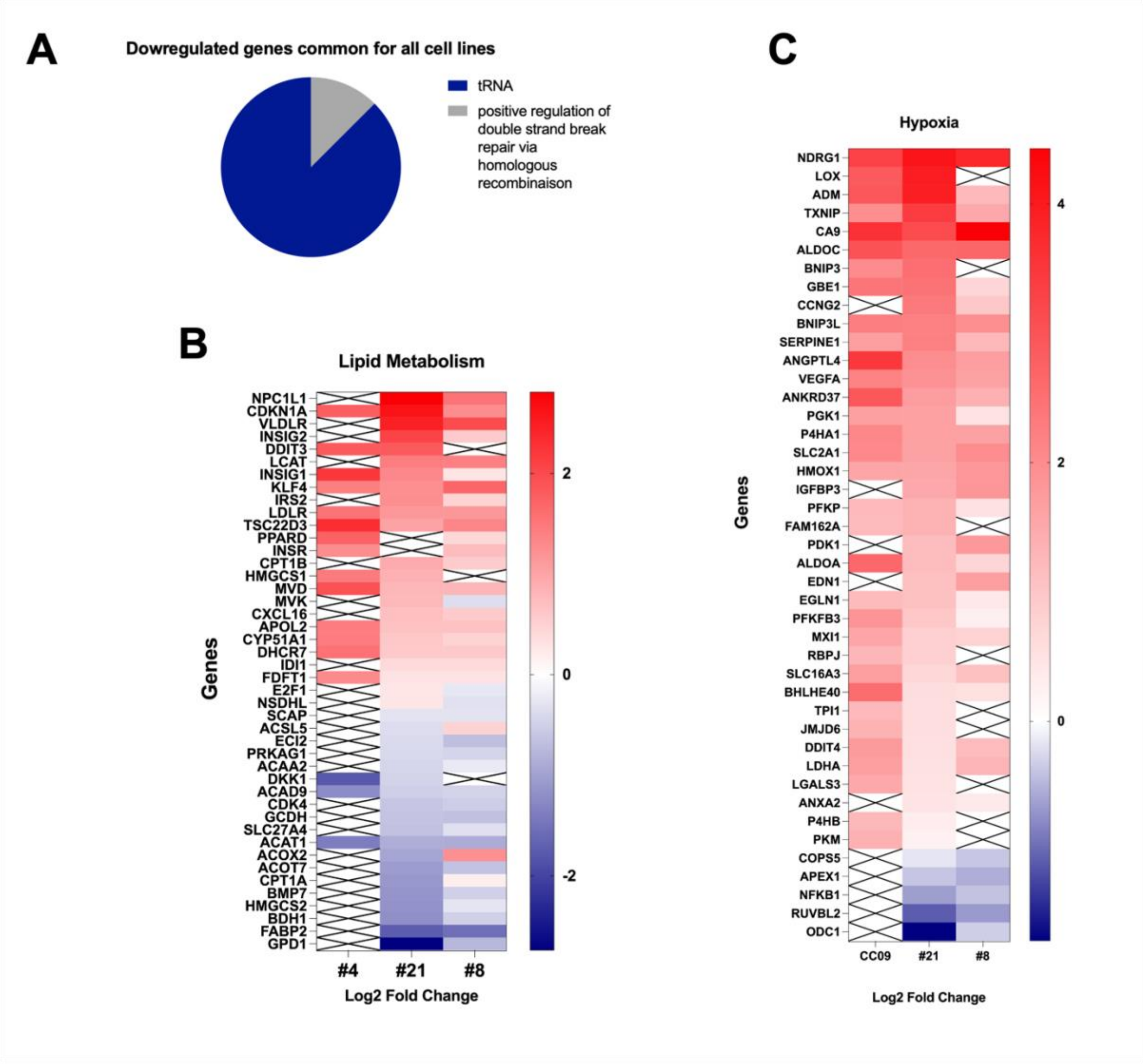


**Figure S5**. **A**) Diagram of common downregulated pathways in all CR-CSCs culture under hypoxia. **B**) Heatmap of proteins belonging to the lipid metabolism pathway. Representation of Log2 Fold change values. **C**) Heatmap of proteins belonging to Hypoxia pathway. Representation of Log2 Fold change values.
